# Antibacterial activity of a newly identified phage-derived endolysin and its parental bacteriophage against clinical uropathogenic *Escherichia coli*

**DOI:** 10.64898/2026.09.09.750462

**Authors:** Wojciech Wesołowski, Sylwia Bloch, Grzegorz Czerwonka, Ernest Jagieła, Łukasz Grabowski, Marta Jeschke, Jakub Jacewicz, Emilia Piwnicka, Joanna Morcinek-Orłowska, Hanna Loika, Paulina Czaplewska, Grzegorz Węgrzyn, Wioletta Adamus-Białek, Aleksandra Łukasiak, Bożena Nejman-Faleńczyk

## Abstract

The increasing prevalence of antibiotic-resistant uropathogenic *Escherichia coli* highlights the need for antibacterial strategies that can complement or extend beyond conventional antibiotic treatment. Bacteriophages and phage-derived lytic enzymes represent promising alternatives because of their distinct mechanisms of bacterial killing and their potential activity against drug-resistant pathogens. In this study, we characterized the newly discovered UPEC-infecting bacteriophage vB-EcoS_57-3 and the endolysin 57_3Lys, encoded by this phage, combining genomic, structural, and functional approaches. The phage demonstrated lytic activity against clinical UPEC isolates and retained antibacterial potential under conditions relevant to the urinary tract. Genomic and sequence analyses revealed distinctive features of 57_3Lys associated with signal-anchor-release endolysins and suggested a less common mode of intracellular translocation and activation. Functional experiments supported the involvement of the bacterial secretion machinery in endolysin-mediated lysis. Notably, the purified enzyme also displayed antibacterial activity against intact clinical *E. coli* cells, despite the intrinsic barrier presented by the Gram-negative cell envelope. Although this activity developed slowly, it significantly reduced both bacterial culture turbidity and viable cell number. Together, these findings provide new insights into the antimicrobial strategies based on the phages and their phage lytic enzymes, supporting further exploration of vB-EcoS_57-3 and 57_3Lys as potential tools against antibiotic-resistant UPEC.

**Highlights:**

- **vB-EcoS_57-3 reveals lytic activity against multiple clinical UPEC isolates.**
- **The phage retains antibacterial activity under artificial urinary conditions.**
- **Endolysin 57_3Lys exhibits distinctive SAR-associated features and activity mechanism.**
- **Purified 57_3Lys lyses intact UPEC cells in the absence of a permeabilizer.**

## 1. Introduction

Urinary tract infections (UTIs) are among the most common bacterial infections worldwide and constitute the second most frequent group of infections after those affecting the respiratory tract. They represent a substantial healthcare burden, affecting approximately 150–250 million people annually and generating considerable medical and socioeconomic costs. Although UTIs occur in both sexes, women are disproportionately affected because of anatomical and physiological predispositions (Salman et al., 2026). Gram-negative bacteria are responsible for the majority of UTIs, with members of the order Enterobacterales, particularly *E. coli* and *Klebsiella pneumoniae*, accounting for most cases (Flores-Mireles et al., 2015; Cito et al., 2026).

Uropathogenic *E. coli* (UPEC) is the predominant etiological agent of UTIs and is responsible for approximately 85% of uncomplicated infections (Terlizzi et al., 2017). The pathogenicity of UPEC is determined by a diverse repertoire of virulence determinants, including adhesins, toxins such as α-hemolysin, biofilm-forming ability, and other factors that facilitate colonization, persistence, and dissemination within the urinary tract (Terlizzi et al., 2017; Fuad et al., 2026). The increasing prevalence of multidrug-resistant UPEC strains has substantially reduced the effectiveness of conventional antibiotic therapy (Khatun et al., 2026). Furthermore, recurrent infections requiring repeated antibiotic administration promote the accumulation and dissemination of antimicrobial-resistance determinants within the urogenital microbiome (Groleau et al., 2026). Consequently, there is an urgent need to develop alternative antibacterial strategies.

Among the most promising alternatives are bacteriophages, viruses that specifically infect bacteria (Łukasiak et al., 2026; Joshi et al., 2026). Phages are the most abundant biological entities on Earth, with an estimated global population of approximately 10^31^ particles, and represent an enormous reservoir of genetic diversity (Hatfull et al., 2015). Their narrow host specificity minimizes disruption of the commensal microbiota, while their ability to replicate at the site of infection enables self-amplifying antibacterial activity (Łukasiak et al., 2026). In parallel with the rapid increase in the number of newly described bacteriophages, both fundamental and translational phage research have expanded considerably over the past decade. Nevertheless, relatively few bacteriophages, particularly those infecting clinically relevant UPEC strains, have been comprehensively characterized, limiting both mechanistic understanding and therapeutic development (Hussein et al., 2025; Łukasiak et al., 2026; Joshi et al., 2026).

Besides whole phages, increasing attention has been directed toward phage-derived lytic enzymes, particularly endolysins, as next-generation antibacterial agents (Bai et al., 2020; Niaz et al., 2025; Wesołowski et al., 2025; Xia et al., 2026). During the lytic cycle, bacterial cell lysis is orchestrated by the coordinated activity of holins and endolysins. Holins permeabilize the cytoplasmic membrane, whereas endolysins hydrolyse glycosidic, peptide, or amide bonds within peptidoglycan, ultimately causing osmotic collapse and rapid bacterial lysis (Young, 2013; Young, 2014).

Currently, phage endolysins are classified into three major classes (Cernooka et al., 2025). Canonical endolysins, the most widespread class, accumulate in the cytoplasm during phage development and gain access to the peptidoglycan only after the formation of large membrane lesions by holins. A distinct group of double-stranded DNA bacteriophages encodes endolysins containing N-terminal signal-anchor-release (SAR) domains (Woźnica et al., 2015). Recently, a third major class of endolysins, termed C-terminal anchor (CTA) endolysins, has been proposed. CTA endolysins possess a C-terminal transmembrane anchor instead of an N-terminal SAR domain and appear to represent a distinct lysis strategy, although their activation mechanism remains to be fully elucidated (Cernooka et al., 2025).

Unlike two other classes, SAR endolysins are exported into the periplasm via the Sec secretion pathway independently of holin-mediated transport. Their SAR domain remains anchored in the inner membrane because it is not cleaved by signal peptidase, whereas the catalytic domain is already localized in the periplasm in an inactive conformation that prevents premature host cell lysis. Subsequent membrane depolarization, typically mediated by pinholins, releases the SAR domain from the membrane, induces a conformational rearrangement of the enzyme, thereby activating its lytic function (Xu et al., 2005; Woźnica et al., 2015). Structural diversity within both the catalytic and SAR domains underlies distinct activation mechanisms, leading to the classification of SAR endolysins into three functional classes (Xu et al., 2004). Despite their unique activation processes and considerable therapeutic potential, only a limited number of SAR endolysins have been experimentally characterized, particularly those encoded by newly isolated bacteriophages infecting clinically important pathogens.

In the present study, we describe a novel UPEC-infecting bacteriophage, vB-EcoS_57-3, together with its SAR endolysin, 57_3Lys. We demonstrate the antibacterial activity of both the phage and its endolysin against clinical multidrug-resistant UPEC isolates. Comparative genomic and functional analyses identify vB-EcoS_57-3 as a newly identified representative of the genus *Veterinaerplatzvirus* within the family *Drexlerviridae* and reveal that 57_3Lys is a class II SAR endolysin whose activity depends on Sec-mediated translocation. These findings expand the current knowledge of SAR endolysins and support their potential application as antibacterial agents against antibiotic-resistant UPEC.

## 2. Materials and methods

### 2.1. Bacterial strains

The *E. coli* strain EC57 and other clinical UPEC isolates: EC02, EC10, EC28, EC47, EC55, EC70, EC106, EC116, EC131, EC149, EC164, EC179, EC296, EC302, and EC353 used in this study, were isolated from patients with urinary tract infections (UTIs), as described previously by Dziuba et al. (2023), and stored at −80°C at the Department of Medical Genetics and Laboratory Diagnostics, Faculty of Medicine, Jan Kochanowski University in Kielce, Poland. Laboratory *E. coli* strains: MG1655, C600, MC1061, BL21 and DH5α were obtained from the strain collection of the Department of Molecular Biology, University of Gdańsk.

### 2.2. Cultivation of bacteria and DNA extraction

All bacterial strains were cultured at 37°C in liquid Luria-Bertani broth (LB; BioShop, Burlington, ON, Canada) or on solid LB medium supplemented with 1.5% agar (LA; BTL, Łódź, Poland). For genomic DNA isolation, bacterial cultures were grown in LB broth with aeration at 200 rpm and 37°C in a shaker incubator. Genomic DNA was extracted using the NucleoSpin Tissue spin column kit (Macherey-Nagel, Düren, Germany) according to the manufacturer’s instructions. DNA concentration and purity were determined spectrophotometrically by measuring absorbance at 260 nm.

### 2.3. Whole-genome sequencing of EC57

The bacterial isolate EC57 was sequenced by a commercial sequencing service using two complementary sequencing approaches: short-read sequencing on the Illumina MiSeq platform and long-read sequencing on the Oxford Nanopore MinION platform to improve genome assembly quality and completeness (Genomed, Warszawa, Poland). Genomic DNA concentration was measured using the PicoGreen reagent (Invitrogen™, Thermo Fisher Scientific, Waltham, Massachusetts, USA). DNA fragmentation was performed by sonication using a Covaris E210 instrument with parameters recommended for Illumina MiSeq library preparation. Libraries for Illumina sequencing were prepared using the NEBNext® Ultra™ II DNA Library Prep Kit for Illumina® (New England Biolabs, Ipswich, Massachusetts, USA) according to the manufacturer’s instructions. Sequencing was performed using paired-end technology (2 × 300 bp) with the MiSeq Reagent Kit v3 (Illumina, San Diego, California, USA). Libraries for Oxford Nanopore sequencing were prepared using the Rapid Barcoding Kit (SQK-RBK004; Oxford Nanopore Technologies, Oxford, UK) and sequenced on a SpotON Flow Cell Mk I (R9.4.1).

### 2.4. De novo assembly and annotation

Raw sequencing reads generated from both sequencing platforms were assembled using a hybrid assembly approach implemented in Unicycler v0.5.0 (Galaxy version 0.5.0+galaxy1) with default parameters. Contigs shorter than 500 bp were discarded. The assembled genome was initially annotated using the Bakta Web server (Schwengers et al., 2021; Beyvers et al., 2025). After submission to NCBI, the genome was reannotated using the NCBI Prokaryotic Genome Annotation Pipeline (PGAP) (Tatusova et al., 2016). The genome sequence of strain EC57 was deposited in GenBank under the whole-genome shotgun (WGS) accession number JAWHTW010000001.1.

### 2.5. Phylogenetic analysis of EC57

A single nucleotide polymorphism (SNP)-based phylogenetic analysis was performed using CSI Phylogeny to reconstruct the phylogenetic tree. The analysis was conducted using the default parameters: minimum depth at SNP positions of ×10, minimum relative depth of 10%, minimum distance between SNPs (prune) of 10 bp, minimum SNP quality score of 30, minimum read mapping quality of 25, minimum Z-score of 1.96, and without exclusion of heterozygous SNPs (Kaas et al., 2014). The resulting Newick-format file was subsequently imported into iTOL for phylogenetic tree visualization and annotation (Letunic and Bork, 2021).

### 2.6. Identification of plasmid replicons, CRISPR-Cas systems, antimicrobial resistance genes, and virulence-associated genes in the EC57 genome

Putative plasmid replicons were identified using PlasmidFinder v3.0.3 (Carattoli et al., 2014). The assembled genome sequence was analyzed using the default parameters, with a minimum nucleotide identity threshold of 95% and a minimum coverage threshold of 60%. Identified replicon sequences were assigned to incompatibility (Inc) groups based on the PlasmidFinder database, enabling characterization of the plasmid content of the EC57 genome.

CRISPR arrays and associated *cas* genes were identified using the CRISPRCasFinder web server (Couvin et al., 2018). The assembled genome sequence was analyzed to identify CRISPR arrays and classify CRISPR-Cas systems based on repeat sequences, spacer content, and the presence of adjacent *cas* genes. CRISPR arrays were assigned confidence levels according to the CRISPRCasFinder scoring system, and CRISPR-Cas subtypes were determined based on the composition and organization of the *cas* operon.

Putative virulence-associated genes (VAGs) were identified using the Virulence Factor Database (VFDB) (Liu et al., 2019). Genes were classified according to their functional categories, including adhesion, invasion, iron acquisition, secretion systems, toxins, autotransporters, and immune evasion.

Antimicrobial resistance genes (ARGs) were identified using the Resistance Gene Identifier (RGI) v6.0.4 against the Comprehensive Antibiotic Resistance Database (CARD) v3.2.9 (Alcock et al., 2023). Protein sequences predicted from the EC57 genome were analyzed using the Perfect and Strict hit criteria with default parameters. Identified ARGs were classified according to their resistance mechanism, AMR gene family, and associated antibiotic class.

### 2.7. Sequencing, assembly and annotation of phage vB-EcoS_57-3

The genome of phage vB-EcoS_57-3 was sequenced by Genomed (Warszawa, Poland) using next-generation sequencing (NGS) on an Illumina MiSeq platform with paired-end technology (2 x 300 bp). A double-stranded DNA (dsDNA) sequencing library was prepared using the NEBNext® Ultra™ II DNA Library Prep Kit (New England Biolabs, Ipswich, Massachusetts, USA). Raw sequencing reads were trimmed using Cutadapt v3.0 (Martin, 2011), and read quality was assessed with FastQC v0.12.1 (https://www.bioinformatics.babraham.ac.uk/projects/fastqc/) by Genomed (Warszawa, Poland). Genome assembly and annotation were performed according to the workflow described by Shen and Millard (2021). *De novo* assembly was carried out using SPAdes v3.15.5 with the “*only-assembler*” option (Bankevich et al., 2012), based on 188,971 sequencing reads. Assembly artifacts were removed using apc.pl (available at: https://github.com/jfass/apc). Viral genome termini, DNA packaging strategy, and genome reordering were predicted using PhageTermVirome v4.1 with Illumina short reads and default parameters (Garneau et al., 2021). The assembled genome was subsequently polished using Pilon v1.24 (Walker et al., 2014) with filtered and sorted reads generated by SAMtools (Danecek et al., 2021). The final assembly consisted of a single contig representing the complete genome of phage vB-EcoS_57-3, with an average sequencing coverage of 1,710x.

Genome annotation was performed using Pharokka v1.5.1 (Bouras et al., 2023). Coding DNA sequences (CDSs) were predicted with PHANOTATE (McNair et al., 2019), whereas tRNA genes were identified using tRNAscan-SE 2.0 (Chan et al., 2021). Functional annotation was generated by comparing predicted CDSs against the PHROGs (Terzian et al., 2021), VFDB (Chen et al., 2005), and CARD (Alcock et al., 2020) databases using MMseqs2 (Steinegger et al., 2017) and PyHMMER (Larralde et al., 2023). Genome visualization, as well as GC content and GC skew analyses, were performed using the Proksee platform (Grant et al., 2023). The phage lifestyle was predicted using PhaTYP (Shang et al., 2023), implemented in PHAbox v2.1.13. To identify antimicrobial resistance determinants and virulence-associated genes within the phage genome, the following tools were used: ABRicate v1.0.1 (Seemann, 2023), AMRFinderPlus v3.11 (Feldgarden et al., 2021), RGI v6.0.3 (Alcock et al., 2023), VirulenceFinder (Joensen et al., 2014), and ResFinder (Bortolaia et al., 2020). Comparative genomic analysis of the related phages CEB_EC3a, DTL, IME542, 12210I, Rtp, IME253, and ACG-M12 was performed using Easyfig v2.2.5 (Sullivan et al., 2011). The assembled and annotated genome sequence of phage vB-EcoS_57-3 was deposited in the GenBank database under accession number PZ349761.

### 2.8. Phylogenetic analysis of vB-EcoS_57-3

To determine the phylogenetic position and taxonomic classification of bacteriophage vB-EcoS_57-3, PhaGCN implemented in PHABOX v2.1.13 (Shang et al., 2021) was initially used to assign the phage to a putative taxonomic family. Subsequently, 211 complete reference genomes representing the identified family were retrieved from the NCBI Virus database. Pairwise intergenomic similarities among all 212 phages, including vB-EcoS_57-3, were calculated using VIRIDIC (Moraru et al., 2020), and the resulting distance matrix is provided in **Supplementary Table S1**. Hierarchical clustering based on complete linkage was performed using the SciPy v1.16.3 library (Virtanen et al., 2020). Orthologous protein groups were identified using OMA v2.6.0 (Altenhoff et al., 2019). Based on the VIRIDIC results, 19 representative phages were selected for multigene phylogenetic analysis. Phages sharing at least 30% genome-wide similarity with vB-EcoS_57-3 were included, provided they did not belong to the same species-level cluster. When multiple phages formed a species-level cluster, a single representative genome was selected using the medoid-based approach. Phage F61 was selected as the outgroup because it represented the most divergent phage while still sharing the orthologous protein set required for multigene phylogenetic analysis, including the endolysin.

Multiple sequence alignments of orthologous protein groups identified by OMA were generated using Clustal Omega v1.2.3 (Sievers and Higgins, 2018) and subsequently trimmed with trimAl v1.4.rev15 (Capella-Gutiérrez et al., 2009) using the *automated1* option. A maximum-likelihood phylogenetic tree was reconstructed from 11 orthologous protein groups (**Supplementary Table S2**) using IQ-TREE v3.1.1 (Wong et al., 2026). The optimal substitution models were selected with ModelFinder Plus (Kalyaanamoorthy et al., 2017), and phylogenetic inference was performed using the edge-linked partition model (Chernomor et al., 2016), with branch support assessed by 1,000 ultrafast bootstrap replicates (Hoang et al., 2018) and 1,000 SH-like approximate likelihood ratio tests (Guindon et al., 2010). The resulting phylogenetic tree was visualized using Dendroscope v3.8.10 (Huson and Scornavacca, 2012).

### 2.9. Isolation of the vB-EcoS_57-3 bacteriophage from urban sewage

Bacteriophage vB-EcoS_57-3 against *E. coli* EC57 was isolated from urban sewage from the Gdańsk Wastewater Treatment Plant in Poland following the protocol described by Jurczak-Kurek et al. (2016), with some modifications. Briefly, 10 mL of the sewage sample was mixed with 1 mL of the host strain overnight culture and cultivated for a few hours at 37°C with shaking. After cell lysis, the sample was centrifuged at 10,000 x *g* for 30 min at 4°C. The obtained supernatant was treated with 4% chloroform (Chempur, Piekary Śląskie, Poland) for 15 min, centrifuged (2,000 x *g*; 10 min; 4°C), and filtered through a 0.22 µm syringe filter with surfactant-free cellulose acetate membrane (SFCA; Thermo Fisher Scientific, Waltham, Massachusetts, USA). In the next step, 10-fold dilutions of the phage lysate were prepared in TM buffer (10 mM Tris-HCl, 10 mM MgSO_4_; pH 7.2). Then, 50 µL of each dilution was mixed with 1 mL of overnight host bacterial culture and 2 mL of top agar. The mixtures were poured onto Petri dishes (Alchem, Toruń, Poland) containing 25 mL of LA medium. After overnight incubation at 37°C, a single phage plaque was transferred to a new flask with EC57 bacterial culture and propagated three times by the procedure described above to obtain purified lysate with vB-EcoS_57-3 particles.

### 2.10. Lysate titration procedure

The titer of the phage lysate was determined using the standard double-overlay method described by Sambrook and Russell (2001), with some modifications. In the first step, the suspension of phage particles was serially 10-fold diluted in TM buffer (10 mM Tris-HCl, 10 mM MgSO_4_; pH 7.2). Then, an appropriate volume of each dilution of phage lysate was mixed with 2 mL of top agar and 1 mL of the overnight bacterial cell culture. The mixtures were poured onto the standard Petri dishes filled with 25 mL of LA. Plates were incubated overnight at 37°C. The phage titer was calculated on the basis of counted plaques and presented as the number of plaque-forming units per milliliter (PFU/mL).

### 2.11. Preparation of the vB-EcoS_57-3 lysate

*E. coli* EC57 was grown in LB medium at 37°C to the early log phase (OD_600_ = 0.2). Then, host bacterial cells were infected with vB-EcoS_57-3 phage at an m.o.i. of 0.05 and cultivated until lysis was observed. The phage suspension was shaken with 4% chloroform (Chempur, Piekary Śląskie, Poland) for 15 min at 37°C, centrifuged at 2,000 x *g* for 20 min at 4°C to separate the host cell debris, and passed through a 0.22 µm syringe filter with SFCA membrane (Thermo Fisher Scientific, Waltham, Massachusetts, USA). The lysate was titrated on double-layer agar plates and incubated overnight at 37°C. The phage titer was determined on the basis of counted plaques. The phage stock was stored at 4°C until subsequent experiments.

### 2.12. Electron microscopy

The lysate with vB-EcoS_57-3 particles was concentrated in the presence of 10% polyethylene glycol 8000 (PEG8000; BioShop, Burlington, ON, Canada) for 18 h at 4°C. After centrifugation (8,000 x *g*; 20 min; 4°C), the supernatant was discarded, and pelleted phage particles were suspended in TM buffer (10 mM Tris-HCl, 10 mM MgSO_4_; pH 7.2). PEG8000 was removed from the sample using 4% chloroform. Concentrated lysate was then subjected to ultracentrifugation (20,000 rpm; 2 h; 10°C) in the cesium chloride gradient as described by Sambrook and Russell (2001). Electron microscopic analysis of vB-EcoS_57-3 virions was performed in the Bioimaging Laboratory, Faculty of Biology, University of Gdańsk, Poland, by using the negative staining with uranyl acetate method as previously described by Topka-Bielecka et al. (2020).

### 2.13. Plaque morphology

The plaque morphology of phage vB-EcoS_57-3 was examined on an overnight culture of the EC57 isolate. To obtain visible plaques, 10-fold dilutions of vB-EcoS_57-3 stock were prepared in TM buffer (10 mM Tris-HCl, 10 mM MgSO_4_; pH 7.2). In the next step, 1 mL of the overnight EC57 culture was mixed with 2 mL of top agar and 15 µL of an appropriate dilution of lysate. The mixture was poured onto a Petri dish with 25 mL of LA medium and incubated overnight at 37°C. The next day, plaque morphology and diameter were determined. Visualization of phage vB-EcoS_57-3 plaques was performed using UVITEC Alliance Q9 Mini software (UVITEC Ltd, Cambridge, UK).

### 2.14. Efficiency of plating (EOP)

Efficiency of plaque formation on 21 *E. coli* strains was determined according to the protocol described by Mirzaei and Nilsson (2015), with some modifications. Briefly, 10-fold serial dilutions of phage stock were prepared in TM buffer (10 mM Tris-HCl, 10 mM MgSO_4_; pH 7.2). Then, 15 µL of the appropriate dilution was mixed with 1 mL of an overnight bacterial culture and 2 mL of top agar. The mixture was poured onto an LA agar plate and incubated overnight at 37°C. Efficiency of plaque formation of vB-EcoS_57-3 phage was calculated by dividing the phage titer on the tested strain by the phage titer on the EC57 host. The EOP values for each virus-bacteria combination were classified as follows: (i) EOP ≥ 0.05 - high infection efficiency, (ii) 0.5 > EOP ≥ 0.1 - medium infection efficiency (iii) 0.1 > EOP ≥ 0.001 - low infection efficiency, and (iv) EOP ≤ 0.001-inefficient infection.

### 2.15. Adsorption assay

A phage adsorption assay was performed following a previously described method by Czajkowski et al. (2014), with slight modifications. Briefly, host bacteria were cultivated to an OD_600_ of 0.2 at 37°C. Then, 3 mL of EC57 cells were mixed with a phage suspension to reach an m.o.i. of 0.05 and incubated at 37°C without shaking. Aliquots from the prepared mixture were collected at 0 (control), 20 s, 1 min, 2 min, 3 min, 4 min, 6 min, 8 min, and 10 min post-infection intervals and centrifuged at 12,000 x *g* for 1 min at 4°C to pellet the host bacteria with adsorbed phages. In the next step, the obtained supernatants were filtered with 0.22 µm syringe filters with SFCA membranes (Thermo Fisher Scientific, Waltham, Massachusetts, USA) and tested for free, non-adsorbed virus particles by using the standard double-overlay method. The kinetics of phage adsorption to the host cell surface was determined based on the formula: percentage of adsorbed phages = ((control titer – residual titer)/control titer) x 100. Additionally, to estimate the number of bacterial cells during the adsorption process, 100 µL of samples were collected at the times indicated above, and their serial 10-fold dilutions were prepared in 0.85% NaCl (STANLAB, Lublin, Poland). Then, 40 µL of each dilution were spread onto LA plates. After overnight incubation at 37°C, the number of bacterial cells per mL (CFU/mL) was calculated.

### 2.16. One-step growth curve

To determine the latent period and the burst size of phage vB-EcoS_57-3, a one-step growth experiment was conducted according to the procedure described by Necel et al. (2020), with some modifications. Host bacteria were grown in LB medium (BioShop, Burlington, ON, Canada) to an OD_600_ of 0.2. Then, 10 mL of bacterial culture was harvested by centrifugation (2,000 x *g*; 10 min; 4°C) and resuspended in 1 mL of LB medium supplemented with 3 mM NaN_3_ (Avantor Performance Materials Poland S.A., Gliwice, Poland) to synchronize host growth. After 5 min incubation of the sample at 37°C, the phage particles were added to bacteria to an m.o.i. of 0.05. The phages were allowed to adsorb to the surface of bacterial cells for 1 min at 37°C. To remove unadsorbed virions, the sample was centrifuged (2,000 x *g*; 10 min; 4°C), and the bacterial pellet was suspended in 1 mL of LB medium supplemented with 3 mM NaN_3_ (Avantor Performance Materials Poland S.A., Gliwice, Poland). In the next step, 25 µL of the suspension was added to 25 mL of LB medium (time 0) and cultivated at 37°C with shaking. The number of infective centers was estimated from the sample taken 1 min after infection by mixing 2 µL of the suspension with 1 mL of the overnight EC57 culture and 2 mL of top agar. Subsequently, the mixture was poured onto an LA plate. At indicated times, all samples were cleared by centrifugation (10,000 x *g*; 2 min; 4°C) and titrated to calculate the value of PFU/mL. After overnight incubation at 37°C, burst size was presented as the ratio of phage titer to the number of infection centers.

### 2.17. Lysis profile

EC57 cells were grown in LB medium to an OD_600_ of 0.2 at 37°C with shaking. Then, phage stock was added to the bacterial host to an m.o.i. of 0.005. A bacterial culture prepared in the same way but without virus supplementation was used as the control. Bacterial growth was monitored by measuring the optical density (OD_600_) every 10 min for up to 70 min. Additionally, the number of bacterial cells per mL (CFU/mL) and phage titer (PFU/mL) were also determined. After overnight incubation at 37°C, bacterial colonies and plaques were counted.

### 2.18. Stability of phage particles in the presence of physical and chemical factors

To determine the effects of various physical and chemical agents on the stability of the phage particles, the procedures described earlier by Jurczak-Kurek et al. (2016) were used, with some modifications. Briefly, 100 µL of phage lysate was mixed with 900 µL of TM buffer (10 mM Tris-HCl, 10 mM MgSO_4_; pH 7.2). Then, the stability of phage suspension was tested in the presence of the following external factors: temperatures (-20°C, 40°C, and 62°C), pH values (2, 4, and 10), organic solvents (chloroform, DMSO, and ethanol), and detergents (CTAB and sarkosyl). Additionally, the effects of osmotic shock on the survival of virus particles were also analyzed. At appropriate times, samples were collected and assayed for phage presence by the standard double-overlay method described above. The percentage of residual infective virions under different conditions was calculated by comparing the initial and post-incubation phage titer (PFU/mL). Phage particles stored in TM buffer at 4°C were used as a control.

### 2.19. Stability and activity of phage virions in artificial urine

Synthetic urine was prepared according to the protocol previously described by Torzewska and Różalski (2015) and consisted of the following components (g/300 mL): 0.195 g of CaCl_2_ x 2H_2_O (BioShop, Burlington, ON, Canada), 0.195 g of MgCl_2_ x 6H_2_O (Chempur, Piekary Śląskie, Poland), 1.38 g of NaCl (STANLAB, Lublin, Poland), 0.69 g of Na_2_SO_4_ (Thermo Fischer Scientific, Waltham, Massachusetts, USA), 0.195 of g sodium citrate (BioShop, Burlington, ON, Canada), 0.006 g of sodium oxalate (Merck KGaA, Darmstadt, Germany), 1.26 g of KH_2_PO_4_ (Avantor Performance Materials Poland S.A., Gliwice, Poland), 0.48 g of KCl (Chempur, Piekary Śląskie, Poland), 0.3 g of NH_4_Cl (Chempur, Piekary Śląskie, Poland), 7.5 g of urea (Avantor Performance Materials Poland S.A., Gliwice, Poland), and 0.33 g of creatine (Thermo Fischer Scientific, Waltham, Massachusetts, USA). The pH of artificial urine was adjusted to values of 4.5, 5.8, and 6.5 by using HCl (Merck KGaA, Darmstadt, Germany) or NaOH (Merck KGaA, Darmstadt, Germany). Then, prepared urines were sterilized by passing through 0.22 µm syringe filters with SFCA membrane (Thermo Fisher Scientific, Waltham, Massachusetts, USA).

In the first part of the experiment, the effect of artificial urine with different pH values on the stability of phage particles was analyzed. Briefly, 100 µL of phage lysate was mixed with 900 uL of synthetic urine with a pH of 4.5, 5.8, or 6.5 and incubated at 37°C. Samples were withdrawn at the indicated times (0, 2, 4, 6, and 8 h), and the concentration of phage particles (PFU/mL) was determined by using double-layer agar plates.

In the second part of the experiment, the activity of phage virions against host bacteria during incubation in the artificial urine with a pH of 4.5 was tested. EC57 bacteria were grown in fresh LB medium to an OD_600_ of 0.2 at 37°C with shaking. In the next step, 3.6 mL of bacterial culture was centrifuged (2,000 x *g*; 10 min; 4°C), and the obtained pellet was suspended in 0.9 mL of the artificial urine with a pH of 4.5. Then, host bacteria were treated with 100 µL of phage lysate with a titer of 3.6 x 10^9^ PFU/mL, corresponding to an m.o.i. of 0.5. Additionally, to assess the stability of bacterial cells in synthetic urine at pH 4.5, 100 µL of TM buffer (10 mM Tris-HCl, 10 mM MgSO₄; pH 7.2) was mixed with 900 µL of the EC57 culture. The mixture was incubated at 37°C for 24 h. Bacterial viability was determined by enumerating colony-forming units (CFU/mL) at 0, 2, 4, 6, 8, and 24 h. At each time point, 100 µL samples were collected, serially diluted 10-fold in 0.85% NaCl (STANLAB, Lublin, Poland), and 40 µL aliquots of each dilution were spread onto LA plates. Following overnight incubation at 37°C, colonies were counted, and the CFU/mL values were calculated.

### 2.20. Cytotoxicity assay

The safety of phage vB-EcoS_57-3 obtained after purification in the cesium chloride gradient was tested with the Thermo Scientific Pierce chromogenic endotoxin quant kit (Thermo Fischer Scientific, Waltham, Massachusetts, USA) and Neutral Red Uptake (NRU) cytotoxicity assay according to the procedure described earlier by Wesołowski et al. (2026), with some modifications. The human bladder cancer cell line (T24) and the human embryonic kidney line (HEK293) were used in the experiment. The NRU cytotoxicity assay was conducted according to the following protocol. T24 and HEK293 cells were grown in McCoy’s 5A Medium (Avantor Performance Materials Poland S.A., Gliwice, Poland) or Dulbecco’s Modified Eagle’s Medium (Gibco, Thermo Fischer Scientific, Waltham, Massachusetts, USA) supplemented with 10% FBS (Thermo Fischer Scientific, Waltham, Massachusetts, USA), 1% penicillin (Thermo Fischer Scientific, Waltham, Massachusetts, USA), and 1% streptomycin (Thermo Fischer Scientific, Waltham, Massachusetts, USA), respectively. Mammalian cells were seeded into 96-well plates at a density of 10^4^ per well and incubated in the presence of 5% CO_2_ at 37°C for 24 h. Then, the culture medium was discarded from each well and cells were treated with 200 µL of a mixture composed of appropriate medium and vB-EcoS_57-3 phage solution with a final titer of 10^8^, 10^9^ or 10^10^ PFU/mL. In addition, 1% Triton X-100 (Merck KGaA, Darmstadt, Germany) and TM buffer (10 mM Tris-HCl, 10 mM MgSO_4_; pH 7.2) were used as positive and negative controls, respectively. Following 24 h incubation (37°C and 5% CO_2_), the mixture was removed, and mammalian cells were washed with 150 µL of 1x PBS (Thermo Fisher Scientific, Waltham, Massachusetts, USA). In the next step, 0.33% NR dye solution (Thermo Fisher Scientific, Waltham, Massachusetts, USA) was added to each well, and the plate was incubated in the dark at 37°C for 45 min. After removal of the NR dye solution, the tested cells were washed with 150 µL of 1x PBS (Thermo Fisher Scientific, Waltham, Massachusetts, USA) and treated with 100 µL of NR desorb solution containing 50% ethanol (Chempur, Piekary Śląskie, Poland) and 1% glacial acetic acid (Merck KGaA, Darmstadt, Germany) in water. The amount of released NR proportional to the amount of viable mammalian cells was measured as A_540_ in the Viroskan microplate reader (Thermo Fisher Scientific, Waltham, Massachusetts, USA). The cell viability was estimated as a percentage of the control values. According to the PN-EN ISO 10993-5:2009 standard, the tested agent is considered cytotoxic if the mammalian cells’ viability is below 70%. Additionally, the stability of vB-EcoS_57-3 particles in the presence of tested cells and media was determined by using the standard double-overlay method and expressed as PFU/mL.

### 2.21. Cloning of the gene encoding the 57_3Lys endolysin

The 57_3Lys gene was cloned from genomic DNA of phage vB-EcoS_57-3 by PCR using Walk DNA polymerase (A&A Biotechnology, Gdańsk, Poland) and specific primers: F_57_3Lys (5’ GCGCCATGGCTCATATGAAGCAGAAGCTGCTAATTA 3’) and R_57_3Lys (5’ TTTAAGCTTCTCGAGACGCTGCAATCCACTTAAACA 3’). The PCR primers were designed by Primer3web (version 4.1.0) and synthesized by Genomed. (Warszawa, Poland). The PCR product was cloned into a commercially available pET26b(+) expression vector (Merck KGaA, Darmstadt, Germany), which carries a C-terminal His-Tag sequence. Following digestion with *Nde*I and *Xho*I restriction endonucleases (Thermo Fisher Scientific, Waltham, Massachusetts, USA), the PCR product was ligated with the *Nde*I-*Xho*I fragment of plasmid pET26b(+) bearing a kanamycin-resistant gene, using the T4 DNA ligase (Thermo Fisher Scientific, Waltham, Massachusetts, USA). The construct pET26b_57_3Lys was introduced into *E. coli* MC1061 by chemical transformation and then verified by sequencing (Genomed, Warszawa, Poland).

### 2.22. In silico endolysin analysis

The Prot-Param module from the Biopython library was used to predict molecular weight, instability index, isoelectric point (pI), and grand average of hydropathy (GRAVY) score (Cock et al., 2009). The aliphatic index was calculated using a previously described formula (Ikai, 1980). A domain search was performed using InterProScan version 5.72-103 (Jones et al., 2014). Additionally, to get deeper insight into the protein structure and its properties, DeepTMHMM (Hallgren et al., 2022), SignalP6.0 (Nielsen, 2025), CD-Search (Wang et al., 2023), and CAMPR3 (Waghu et al., 2015) were applied.

### 2.23. Overproduction and purification of the 57_3Lys endolysin

The construct pET26b_57_3Lys was transformed into the BL21-AI^TM^ *E. coli* strain (Thermo Fisher Scientific, Waltham, Massachusetts, USA). Bacteria carrying the overexpression plasmid were cultivated at 37°C in 200 mL of LB medium supplemented with 30 µg/mL kanamycin (BioShop, Burlington, ON, Canada) and 12.5 µg/mL tetracycline (Merck KGaA, Darmstadt, Germany) to an OD_600_ of 0.8. At that point, overproduction of the 57_3Lys protein was induced by adding 0.5 mM isopropyl-β-D-thiogalactopyranoside (IPTG; A&A Biotechnology, Gdańsk, Poland) and 0.2% L-(+)-arabinose (BioShop, Burlington, ON, Canada), and the incubation was continued for 1 h at 37°C until complete lysis of bacterial cells occurred. Then, Pierce Protease Inhibitor Tablets (Thermo Fisher Scientific, Waltham, Massachusetts, USA) were added to the lysate and the mixture was shaken for 30 min at 37°C. Subsequently, the lysate was clarified by centrifugation (6,000 x *g*; 10 min; 4°C) and filtered through a 0.45 µm vacuum filter with a polyethersulfone membrane (PES; Avantor Performance Materials Poland S.A., Gliwice, Poland). In the next step, the lysate was loaded onto a 5 mL HisTrap^TM^FF crude column (Cytiva, Marlborough, Massachusetts, USA). The protein was eluted with elution buffer (50 mM Tris-HCl, pH 7.5; 300 mM NaCl, 5% glycerol, 0.2 mM TCEP, and 200 mM imidazole) and dialyzed overnight against buffer containing 50 mM Tris-HCl (pH 7.5; BioShop, Burlington, ON, Canada), 200 mM NaCl (STANLAB, Lublin, Poland), 40% glycerol (STANLAB, Lublin, Poland), and 0.2 mM TCEP (Merck KGaA, Darmstadt, Germany). The purity of the 57_3Lys endolysin was assessed with the use of 20% sodium dodecyl sulfate-polyacrylamide gel electrophoresis (SDS-PAGE), and the concentration was determined fluorometrically by the Qubit™ Protein Assay Kit (Thermo Fisher Scientific, Waltham, Massachusetts, USA). Afterwards, the identity of the purified protein was further verified by LC-MS/MS analysis of excised and in-gel digested protein bands, as described previously (Fiołka et al., 2021), using an M5 MicroLC system with a C18 column coupled to a ZenoTOF 7600 mass spectrometer (SCIEX). The purified protein was stored at −20°C until further analysis.

### 2.24. Sec machinery assay

To test whether Sec machinery is responsible for endolysin secretion, the host cell lysis kinetics by 57_3Lys was determined in the presence of various concentrations of NaN_3_ according to the protocol described by Bai et al. (2020), with some modifications. Briefly, *E. coli* BL21-AI^TM^ cells bearing plasmid pET26b_57_3Lys were grown in LB medium at 37°C to an OD_600_ of 0.8 with aeration. Then, overproduction of 57_3Lys from the pET26b_57_3Lys plasmid in host cells was induced by 0.5 mM IPTG (A&A Biotechnology, Gdańsk, Poland) and 0.2% L-(+)-arabinose (BioShop, Burlington, ON, Canada). Additionally, some cultures were treated with NaN_3_ added to a final concentration of 1 mM, 5 mM, or 10 mM. Controls were grown without addition of induction agents and NaN_3_ or in the presence of 10 mM NaN_3_. Bacteria were incubated at 37°C with shaking, and their density was monitored by OD_600_ measurement at the same time intervals, every 10 min for 180 min.

### 2.25. Activity assay of the 57_3Lys endolysin

Turbidity and cell number reduction assays of EC47, EC57, and EC131 cultures by purified 57_3Lys endolysin were performed at 37°C in a standard 96-well titration plate. Briefly, tested bacteria were grown in LB medium (BioShop, Burlington, ON, Canada) at 37°C to an OD_600_ of 0.8. Then, 50 mL of bacterial culture was spun down (2,000 x *g*; 20 min; RT) and subsequently suspended in 5 mL of 50 mM Tris-HCl (pH 7.5). Bacteria were shaken for 45 min at RT. To measure the protein activity, 200 µL of 57_3Lys endolysin at a concentration of 60 ng/µL or 200 µL of the reaction buffer (negative control; 50 mM Tris-HCl, pH 7.5; 200 mM NaCl; 40% glycerol, and 0.2 mM TCEP) were mixed with 100 µL of tested bacterial culture. Additionally, the antibacterial activity of the chicken egg white lysozyme (Merck KGaA, Darmstadt, Germany) against host bacteria was determined. The changes in OD_600_ were recorded after 24 h of incubation at 37°C by using the Viroskan microplate reader (Thermo Fisher Scientific, Waltham, Massachusetts, USA). During this experiment, the number of survivors per mL was also examined. To estimate CFU/mL, 20 µL of bacterial samples were collected and 10-fold diluted in 0.85% NaCl (STANLAB, Lublin, Poland). In the next step, 10 µL of each dilution was spotted onto a LA plate. After overnight incubation at 37°C, colonies were counted, and the CFU/mL was calculated.

### 2.26. Statistical analysis

All experiments were carried out in at least three or even more biological replicates. The means ± SD were used in the statistical analysis of the data and the graphics. A comparison of two averages in experiments was performed using Student’s *t*-test by using Microsoft^®^ Excel 365 with Real Statistics Resource Pack. The significant differences were marked by asterisks as follows: *p* < 0.01 (**) or *p* < 0.001 (***).

## 3. Results

### 3.1. Overview of the uropathogenic clinical EC57 isolate, a primary host for phage vB-EcoS_57-3

#### 3.1.1. EC57 genomic content and pathogenicity determinants

*E. coli* EC57 was part of the clinical *E. coli* strain collection previously analyzed by Dziuba et al. (2023). This strain was isolated from the urine of a 14-year-old boy and belongs to the B2 phylogenetic group.

According to Bakta annotation, the genome of *E. coli* EC57 consists of one complete circular chromosome and three plasmids with a total size of 5,111,555 bp and was assembled into a single large chromosomal contig. Genome annotation performed using Bakta identified a total of 5,108 genomic features, including 4,684 protein-coding sequences (CDSs) and 65 small open reading frames (sORFs). In addition, 88 tRNA genes, 22 rRNA genes, one tmRNA, 189 ncRNAs, and 52 ncRNA-associated regions were identified. The relatively high number of regulatory ncRNAs suggests complex post-transcriptional regulation and adaptive potential under changing environmental conditions. The genome also contained two predicted chromosomal origins of replication (*oriC*), as well as one plasmid replication origin (*oriV*) and one origin of transfer (*oriT*), confirming the mobilizable nature of at least one plasmid present in the strain. Overall, the genome architecture of *E. coli* EC57 reflects a genetically versatile strain with a well-conserved core genome supplemented by plasmid-associated functions that may contribute to environmental adaptation, survival, and acquisition of accessory traits, including antimicrobial resistance or virulence-associated determinants.

PlasmidFinder tool identified a single plasmid, designated pEC57-1, in the *E. coli* EC57 genome. The plasmid DNA consists of 69,890 bp, and pEC57-1 was classified as a member of the IncFII incompatibility group, showing 100% identity to the IncFII replication region. Comparative analysis revealed that the replication region of plasmid pEC57-1 is identical to that of the *E. coli* plasmid pC15-1a (ENA accession number AY458016), showing 258/261 bp coverage and 100% nucleotide identity according to PlasmidFinder. However, comparison of the complete plasmid sequence using BLAST indicated that pEC57-1 is most closely related to plasmid p1_A28403 from *Enterobacter hormaechei* (accession number CP182042.1), with approximately 90% query coverage and 99.74% nucleotide identity. These results suggest that although pEC57-1 carries an IncFII replication region identical to that of pC15-1a, its overall sequence differs substantially from previously described plasmids. Even after lowering the detection thresholds to 60% minimum identity and 30% minimum coverage, no additional plasmid replicons were identified. However, further examination of the genome assembly revealed two additional circular sequences potentially representing plasmids, they contained no recognizable plasmid replicon.

CRISPRCasFinder identified eight CRISPR loci in the *E. coli* EC57 genome (**Table 1**). Six loci (EC57-1, EC57-2, EC57-3, EC57-6, EC57-7, and EC57-8) consisted of single-spacer CRISPR arrays lacking adjacent *cas* genes and were classified as low-confidence structures (evidence level 1). A complete, high-confidence Type I-F CRISPR-Cas system (evidence level 4) was identified between positions 3,151,428 and 3,162,042 bp. This locus comprised two CRISPR arrays (EC57-4 and EC57-5) containing 15 and 10 spacers, respectively, flanking a complete *cas* operon encoding *cas6*, *csy3*, *csy2*, *csy1*, *cas3-cas2*, and *cas1*. Both arrays shared the repeat consensus sequence TTTCTAAGCTGCCTGTACGGCAGTGAAC and were oriented in the reverse direction (**Table 1**).

**Table 1.**
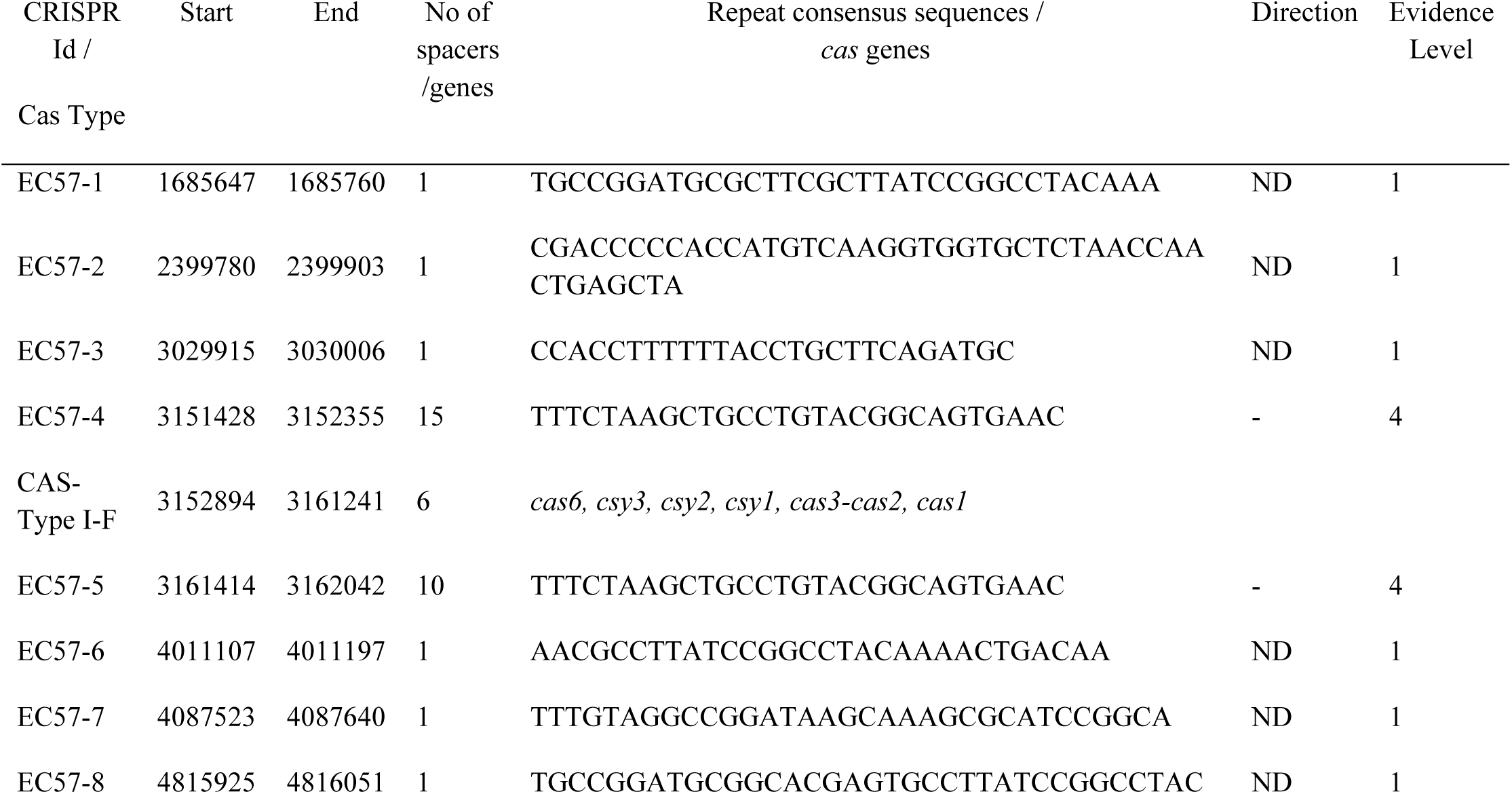
CRISPR arrays and *cas* gene sequences detected in EC57 genome.

CARD analysis identified two acquired antimicrobial resistance genes, *blaTEM-1* and *tet(A)*, encoding a TEM-1 β-lactamase and a tetracycline efflux pump, respectively. In addition, several chromosomally encoded antimicrobial resistance determinants were detected, including multidrug efflux systems (*acrAB-tolC, acrDEF, mdtABC-EFGHMNOP*, and *emrABKYR*) and regulatory systems associated with antimicrobial resistance (*evgAS*, *baeRS*, *marA*, *soxRS*, and *crp*). The genome also contained genes implicated in resistance to peptide antibiotics (*arnT*, *pmrF*, *ugd*, and *eptA*) and β-lactam resistance (*ampC*). Furthermore, CARD identified resistance-associated protein variants, including PBP3 encoded by *ftsI* and GlpT encoded by *glpT*.

VFDB analysis identified a broad repertoire of virulence-associated genes (VAGs) in *E. coli* EC57. The genome contained multiple adhesin-associated genes, including the complete *E. coli* common pilus (*ecp*) and type 1 fimbrial (*fim*) operons, as well as genes associated with F1C (*foc*), P (*pap*), and S (*sfa*) fimbriae, hemorrhagic *E. coli* pilus (*hcp*), the EaeH adhesin (*eaeH*), and UpaG (*upaG*). Several genes encoding autotransporter proteins were also detected, including *agn43*, *cah*, *cdiA*, *ehaB*, *pic*, *upaH*, and *vat*.

The genome also contained invasion-associated genes (*ibeABC* and *tia*) and genes involved in multiple iron acquisition systems, including the *chu* heme uptake system, the salmochelin *iro* operon, and the yersiniabactin-associated genes *fyuA*, *irp1*, *irp2*, and *ybtAEPQSTUX*. Furthermore, genes encoding several toxins were identified, including α-hemolysin (*hlyABCD*), cytotoxic necrotizing factor 1 (*cnf1*), uropathogenic-specific protein (*usp*), and hemolysin E/cytolysin A (*hlyE/clyA*). Genes encoding components of type VI secretion systems (T6SSs), including the ACE-T6SS and SCI-I T6SS, were also detected, together with *espL1*, which encodes a non-LEE-encoded type III secretion system effector.

#### 3.1.2. EC57 phylogeny

SNP-based phylogenetic analysis demonstrated that *E. coli* EC57 clustered within the uropathogenic *E. coli* (UPEC) lineage and was clearly separated from the reference strain *E. coli* K-12 MG1655 (**Figure 1**). The K-12 reference strain grouped together with strains VR50, EcPF16, CI5, and EcPF15, forming a distinct branch distant from EC57. In contrast, EC57 formed a well-supported subclade (bootstrap support = 1.000) with strains 04-00955 and VG38-53E1, while strain A4 represented the closest sister lineage to this cluster (bootstrap support = 0.989). The short branch lengths separating EC57 from 04-00955 and VG38-53E1 indicate a high degree of genomic similarity and recent common ancestry. The EC57-containing clade was clearly separated from other well-characterized UPEC reference strains, including CFT073, UTI89, J96, and 536, which formed independent phylogenetic lineages. These results confirm that EC57 belongs to the UPEC phylogenetic group while representing a distinct evolutionary lineage within this pathotype.

**Figure 1.**
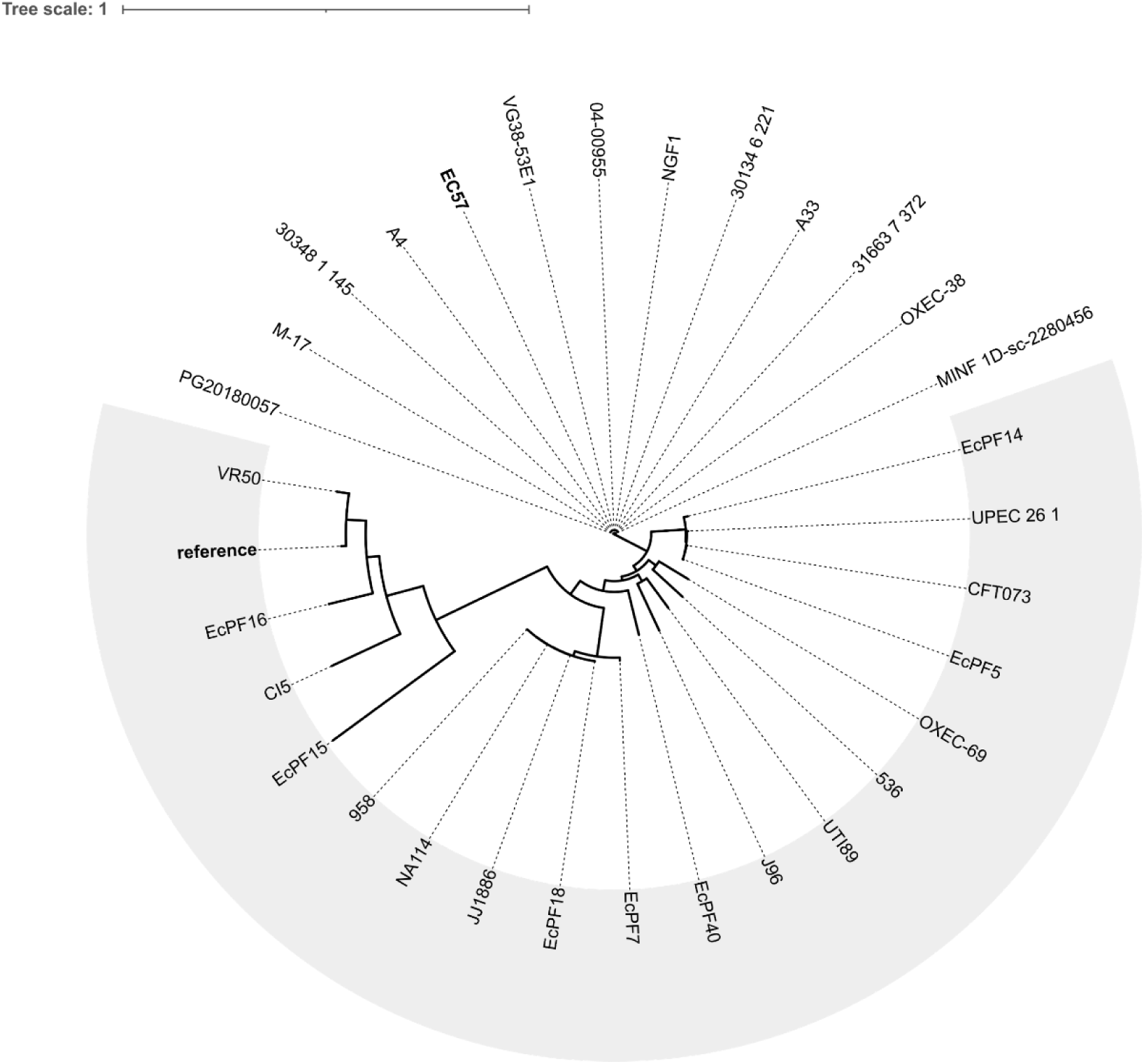
SNP-based phylogenetic tree of *E. coli* EC57 and representative *E. coli* strains. The phylogenetic tree was constructed using whole-genome single nucleotide polymorphism (SNP) analysis performed with CSI Phylogeny and visualized using iTOL. The laboratory strain *E. coli* K-12 MG1655 (reference) was included as a non-pathogenic reference genome. The grey-shaded sector indicates reference UPEC strains included for phylogenetic comparison.

### 3.2. Phage vB-EcoS_57-3 isolation and morphological characterization

The above-characterized UPEC strain EC57 was used to isolate vB-EcoS_57-3 bacteriophage from samples of urban sewage. The isolation procedure was based on the one-host enrichment method in which a raw urban sewage sample was mixed with a culture of the EC57 strain to obtain the lysate of vB-EcoS_57-3 bacteriophage, as described in the Materials and Methods section. The bacteriophage vB-EcoS_57-3 was morphologically characterized using transmission electron microscopy. The virion exhibited a typical siphovirus-like morphology with an icosahedral capsid and a long, flexible, non-contractile tail (**Figure 2A**). Large plaques (diameter 4.5 ± 0.5 mm), with a halo, formed on the *E. coli* EC57 lawn, confirmed its lytic activity against the EC57 isolate and suggested the production of free-releasing polysaccharide depolymerase protein (**Figure 2B**).

**Figure 2.**
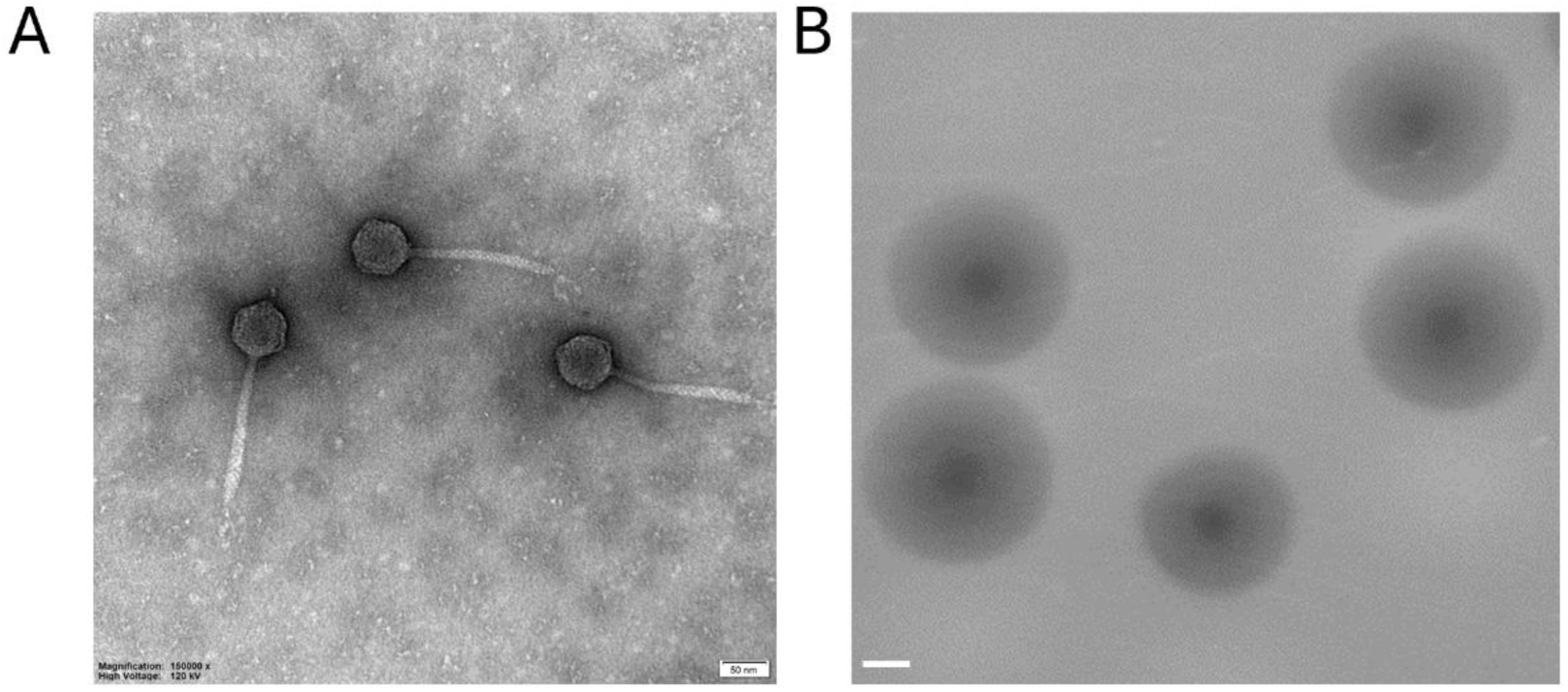
Electron micrograph of phage vB-EcoS_57-3 (**A**) and its plaques formed in double-layer agar plates with *E. coli* EC57 strain (**B**). The bar corresponds to 50 nm on panel A and 1 mm on panel B.

### 3.3. Genome content

The vB-EcoS_57-3 phage possesses a 45,774 bp genome with 91 predicted ORFs, and a GC content of 44%. The whole genome of vB-EcoS_57-3 has been sequenced and deposited in GenBank (accession number PZ349761). Genome annotation revealed modular organization, including genes associated with, e.g., DNA replication, transcription, virion structure, host interaction, and lysis (**Figure 3**). Among all identified coding regions, only 31 ORFs were predicted to be functional genes. Phage lytic enzymes, holin and endolysin, have been identified automatically during annotation by Pharokka, while phage polysaccharide depolymerase was first recognized as a structural protein and subsequently classified by the Phyre2 program as Latka et al., (2019) proposed. Interestingly, only a single (arginine) tRNA gene was predicted in the phage genome (**Figure 3**). In addition, analysis using PhageTermVirome indicated that vB-EcoS_57-3 possesses Headful packing, whereas the PhaTYP classified it as virulent with a confidence equal 1.0. Finally, the comparative genomic analysis (**Figure 4**) performed by the EasyFig program showed conserved genome organization of vB-EcoS_57-3 with closely related phages CEB_EC3a, DTL, IME542, 12210I, Rtp, IME253, and ACG-M12.

**Figure 3.**
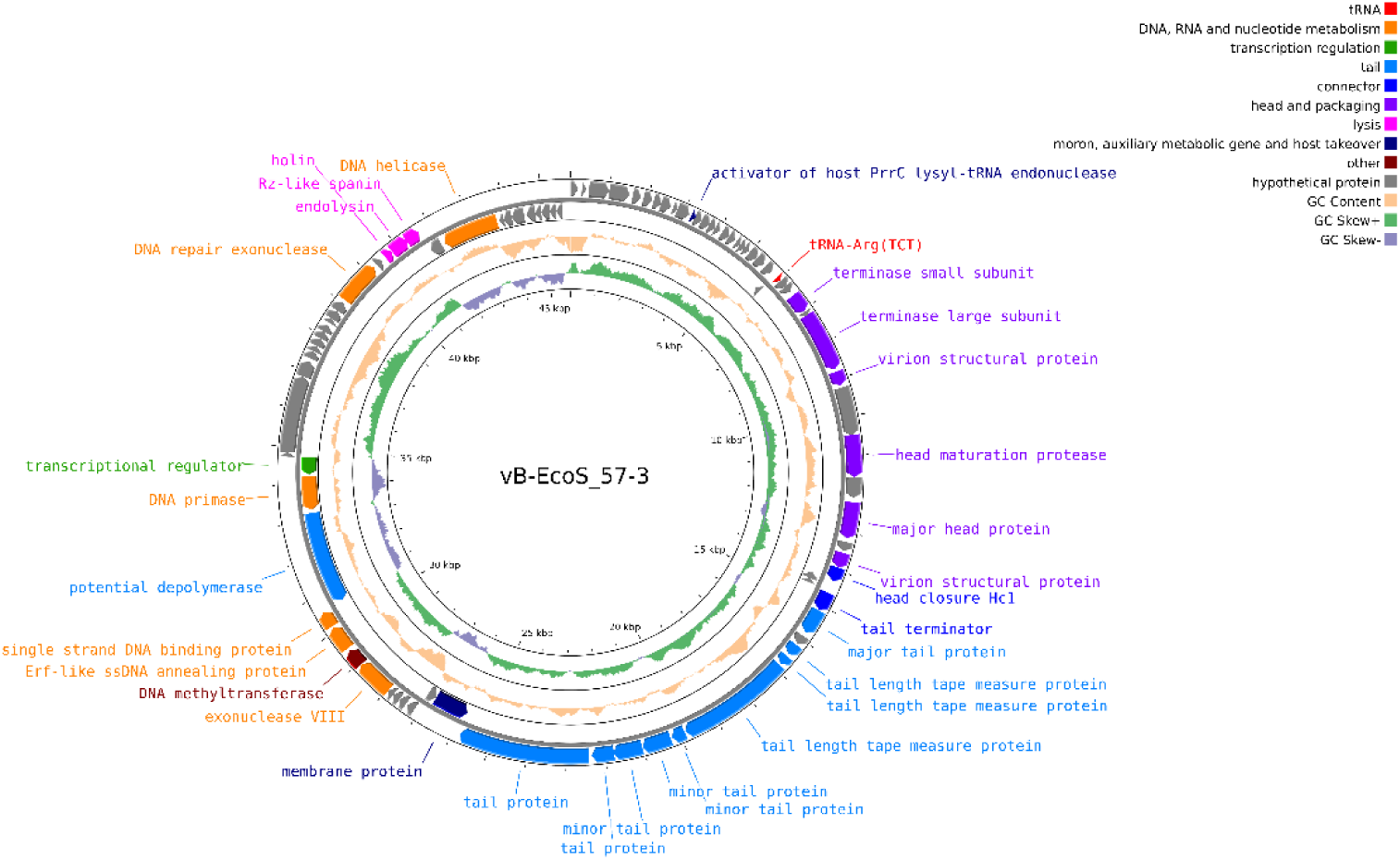
Genome map of bacteriophage vB-EcoS_57-3. Predicted open reading frames (ORFs) are displayed as arrows indicating transcriptional orientation and are color-coded according to functional categories, including DNA, RNA and nucleotide metabolism, transcription regulation, tail, connector, head and packaging, lysis, integration and excision, moron/auxiliary metabolism and host takeover, other functions, and hypothetical proteins (legend on the right side). The inner rings represent GC content and GC skew across each genome. Annotated genes involved in genome replication, virion structure and assembly, host interaction, and lysis are indicated. Genome coordinates (kbp) are shown inside each map.

**Figure 4.**
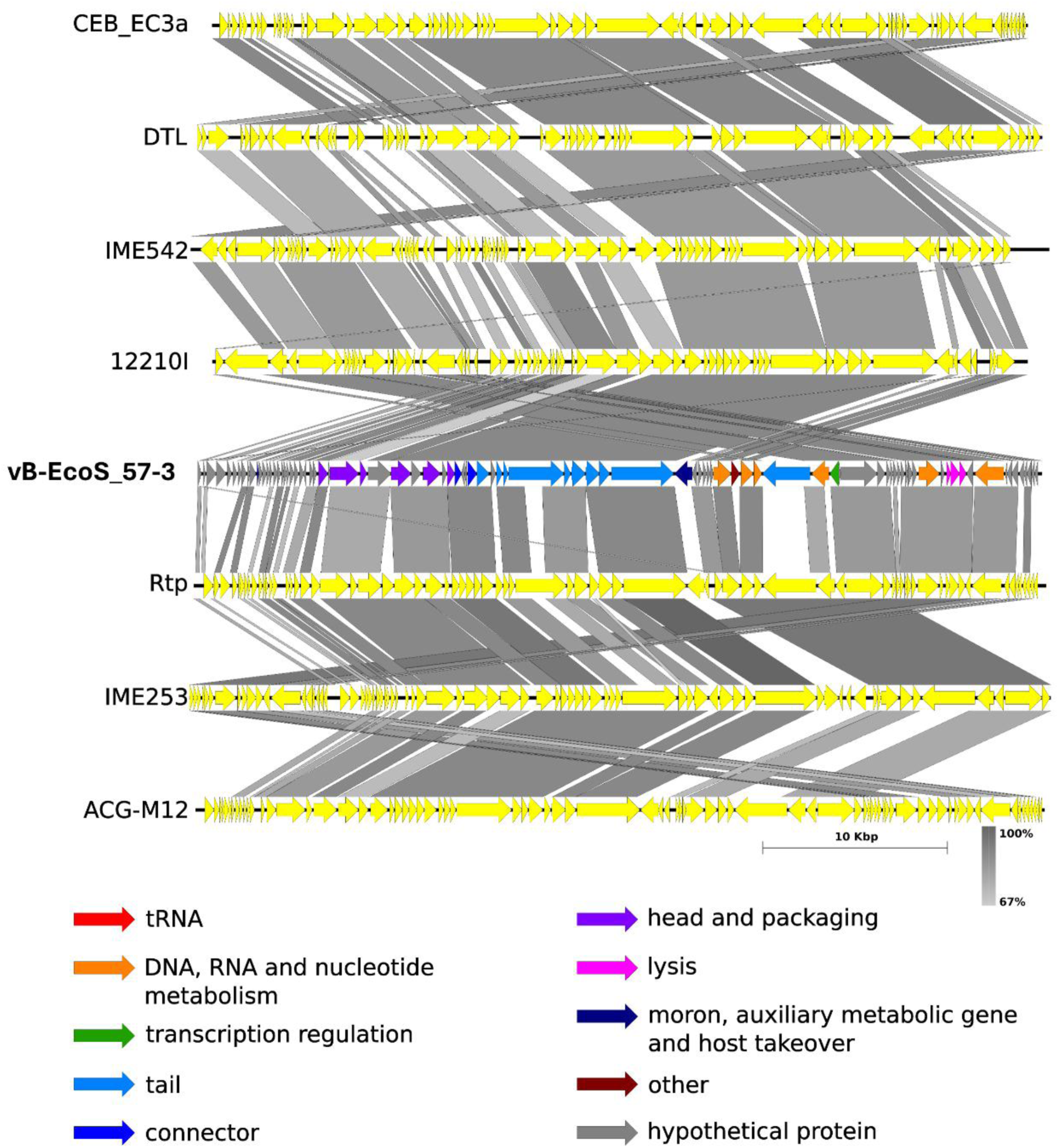
Comparative genome organization of vB-EcoS_57-3 linear genome bacteriophage with their closest related reference phages. Predicted open reading frames (ORFs) are displayed as arrows indicating gene orientation and are color-coded according to functional categories listed in the legend. Grey-shaded regions between genomes represent nucleotide sequence similarity, illustrating conserved synteny and homologous genomic regions. The scale bar represents the percentage of nucleotide sequence identity. The comparison highlights both conserved genome architecture within related phages and structural divergence, supporting the taxonomic distinction of the newly characterized phages. Comparisons were made using Easyfig software.

### 3.4. Taxonomic and phylogenetic analyses

According to the PhaGCN analysis, phage vB-EcoS_57-3 was classified within the family *Drexlerviridae* and the subfamily *Braunvirinae*, with confidence scores of 0.99 and 0.61, respectively. To further investigate its taxonomic position, VIRIDIC was used to assess intergenomic relationships between vB-EcoS_57-3 and *Drexlerviridae* reference phages available in RefSeq (**Supplementary Figure S1 and Supplementary Table S1**). VIRIDIC analysis showed that vB-EcoS_57-3 and vB_EcoS-12210I share 71.45% intergenomic similarity (**Supplementary Table S1**), supporting the assignment of vB-EcoS_57-3 to the genus *Veterinaerplatzvirus* (**Supplementary Figure S1**).

Phylogenetic analysis of 20 phages based on the concatenated alignments of 11 orthologous protein groups (**Supplementary Table S2**) placed vB-EcoS_57-3 within a well-supported clade comprising the *Braunvirinae* phages CEB_EC3a, DTL, IME542, vB_EcoS-12210I, Rtp, IME253, and ACG-M12 (**Figure 5**). Within this clade, vB-EcoS_57-3 was most closely related to vB_EcoS-12210I, with the two phages forming a strongly supported sister group (99.8/95), further supporting their close evolutionary relationship. The 11 orthologous protein groups used for phylogenetic reconstruction included DNA primase, major head protein, endolysin, a single-stranded DNA-binding protein, a virion structural protein, several tail-associated proteins, and one hypothetical protein (**Supplementary Table S2**).

**Figure 5.**
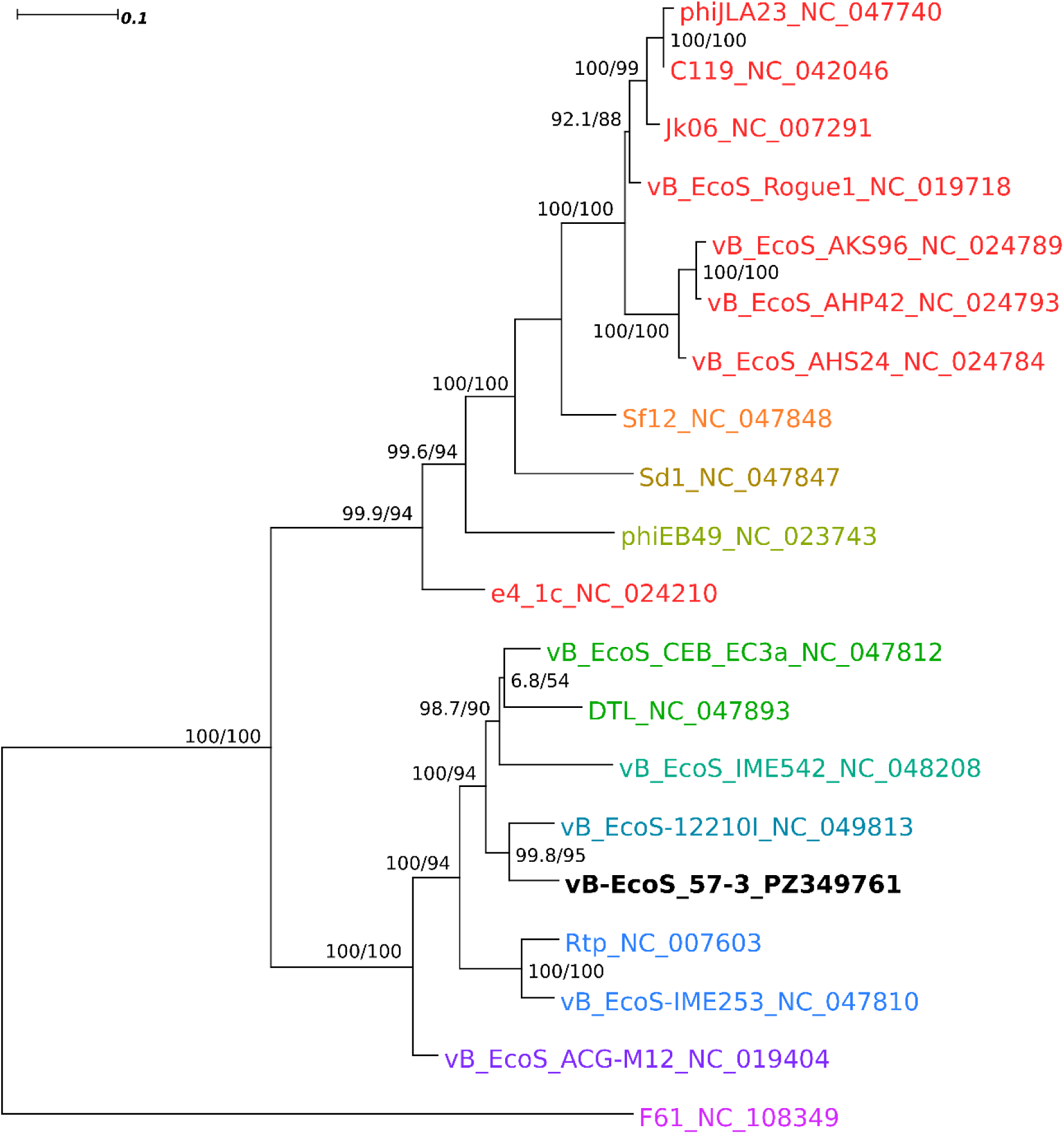
Maximum-likelihood phylogenetic tree of bacteriophage vB-EcoS_57-3 and related phages based on orthologous protein sequences. The phylogenetic tree was reconstructed from concatenated alignments of 11 orthologous protein groups shared among 20 phages (**Supplementary Table S2**) using IQ-TREE v3.1.1. Phylogenetic inference was performed using an edge-linked partition model, with the optimal substitution model for each partition selected by ModelFinder Plus. Branch support values are shown at the nodes as SH-aLRT support (%) / ultrafast bootstrap support (%), based on 1,000 replicates each. Phage vB-EcoS_57-3 is highlighted in bold. Phage F61 was used as the outgroup. Different name colors indicate the genus-level taxonomic assignment of the tested phages as follows: red *Rogunavirus*, orange *Eastlansingvirus*, dark yellow *Wilsonroadvirus*, bottle green *Lindendrivevirus*, green *Loudonvirus*, teal *Christensenvirus*, dark cyan *Veterinaerplatzvirus*, blue *Rtpvirus*, violet *Guelphvirus*, pink *Tlsvirus*. The scale bar represents the number of amino acid substitutions per site.

### 3.5. Host range and specificity against uropathogens

Host range analysis (**Table 2**) indicated variable infectivity efficiency across tested *E. coli* strains. Interestingly, phage vB-EcoS_57-3 infects only UPEC bacteria and shows no lytic activity against non-pathogenic laboratory *E. coli* strains. Among tested uropathogens (Dziuba et al., 2023), high or medium efficiency of plating (EOP ≥ 0.5 or 0.1, respectively) was observed for six clinical isolates, including EC28, EC47, EC57, EC116, EC131, and EC353, whereas the other ten strains showed low, limited, or no susceptibility (**Table 2**). They were previously characterized based on antimicrobial susceptibility and virulence potential (Dziuba et al., 2023). Half of them were susceptible to all antibiotics, while the remaining showed variable resistance to penicillins, cephalosporins, aminoglycosides, or fluoroquinolones. All were susceptible to carbapenems and were ESBL-negative. Besides, most of the tested strains (72%) belonged to the B2 phylogenetic group, all revealed CRISPR-Cas regions, and all (except one) possessed virulence-associated genes (Dziuba et al., 2023) and the ability to form biofilms (data not shown). Therefore, we conclude that this phage is a moderate host-range virus, however, specific to uropathogenic *E. coli* bacteria.

**Table 2.** EOP (efficiency of plating) analysis of vB-EcoS_57-3 phage on *E. coli* laboratory strains and UPEC clinical isolates.

|  | *E. coli*<br>strain | EOP after<br>vB-EcoS_57-3<br>infection* |
| --- | --- | --- |
| **Laboratory *E. coli* strains** | MG1655 | 0 |
|  | C600 | 0 |
|  | MC1061 | 0 |
|  | BL21 | 0 |
| | DH5α | 0 |
| **Clinical UPEC isolates** | EC02 | 0 |
| | EC10 | $\leq 0.001$ |
|  | EC28 | 1.48 |
|  | EC47 | 1.37 |
|  | EC55 | 0 |
|  | EC57 | 1 |
|  | EC70 | 0.007 |
| | EC106 | $\leq 0.001$ |
|  | EC116 | 0.72 |
|  | EC131 | 0.42 |
|  | EC149 | 0 |
|  | EC164 | 0 |
| | EC179 | $\leq 0.001$ |
|  | EC296 | 0 |
|  | EC302 | 0 |
|  | EC353 | 0.84 |

\*According to the accepted classification proposed by Mirzaei et al. (2015) an EOP value $\leq 0.001$ indicates sporadic production of phage particles in a given bacterial strain, while values in the range $0.001 < \text{EOP} < 0.1$ ; $0.1 \leq \text{EOP} < 0.5$ , and $\geq 0.5$ indicate low, medium, and high efficiency of phage infection, respectively. Lack of infection is indicated by 0. The obtained results are from three independent replicates. The SD values are not shown; however, they do not exceed 15%.

### 3.6. Bacteriophage development

The parameters of phage vB-EcoS_57-3 development were determined in the EC57 host strain. The analyzed lysis profile of infection (**Figure 6**) revealed a rapid decrease in OD_600_ (**Figure 6A**) and viable cell counts up to 100 cells in 30 minutes after addition of phage (**Figure 6B**). These drops were accompanied by a significant increase in phage particles up to 10^10^ PFU/mL (**Figure 6C**), confirming efficient lytic development. One-step growth analysis (**Figure 6D**) demonstrated efficient phage replication, with a burst size of approximately 42 PFU per cell and an 11-minute-long latent period.

**Figure 6.**
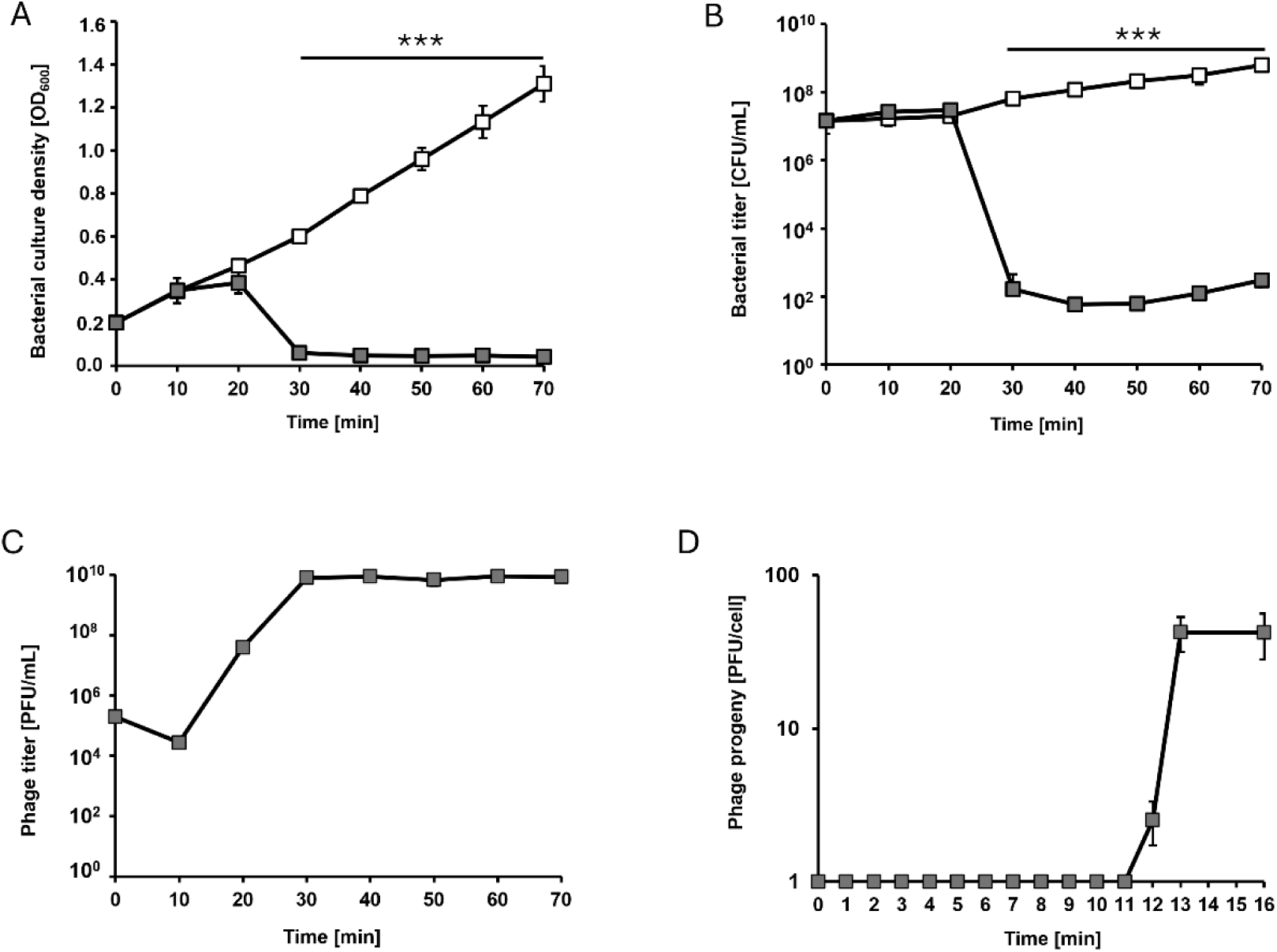
Kinetics of lytic development of bacteriophage vB-EcoS_57-3 in *E. coli* EC57 bacteria (**A-C**) and progeny production by vB-EcoS_57-3 phage in the single life cycle (**D**). Filled squares represent the bacterial strain EC57 following infection with phage vB-EcoS_57-3, whereas open squares represent the uninfected control EC57 without phage addition. Results are shown as (**A**) bacterial culture density measured at OD_600_, (**B**) a number of surviving cells after the vB-EcoS_57-3 infection per 1 mL (CFU/mL), (**C**) a number of phages per 1 mL (PFU/mL), and (**D**) PFU per 1 cell. The average burst size in EC57 at the 16^th^ minute after infection has been estimated to be 42 PFU per cell, indicating efficient lytic development of the phage. Results are presented as mean values ± SD from three independent experiments. Please note that in some cases, the bars are smaller than the sizes of the symbols. The Student’s *t*-test was used to calculate statistical significance. The significance of differences between infected cells and control EC57 culture is observed and marked by asterisks: *P* < 0.001 (***).

Adsorption of vB-EcoS_57-3 on cells of the EC57 host (**Figure 7A**) was moderately fast but efficient. Initially, a rapid rise in adsorption efficiency was observed, suggesting a fast initial interaction phase between phage particles and bacterial cells, however, this was followed by a transient decrease in the percentage of adsorbed phages, which may reflect short-term binding instability. From the second minute, the adsorption efficiency increased again and continued to rise more gradually, approaching a plateau efficiency equal to 80% at the 10^th^ minute, indicating a possible saturation of available binding sites on the bacterial surface. Importantly, bacterial cell numbers remained stable during all analyzed adsorption stages (**Figure 7B**), proving that the observed drop was not an effect of phage entry inside bacteria and its further lytic development leading to the decrease in bacteria number, but likely due to the reversible phage binding to the cell surface at the early stages of the infection.

**Figure 7.**
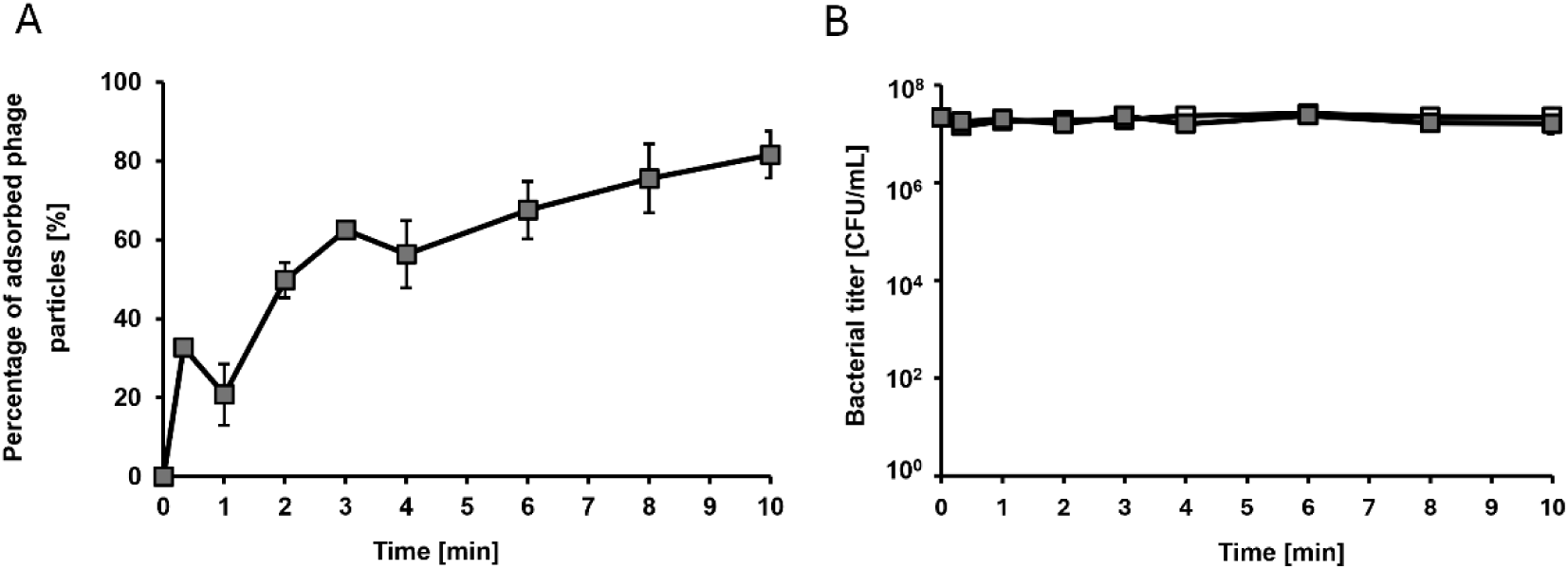
Kinetics of vB-EcoS_57-3 phage adsorption on host EC57 cells (**A**) and number of bacterial cells during this process (**B**). Filled squares represent the bacterial strain EC57 following infection with phage vB-EcoS_57-3, whereas open squares represent the uninfected control EC57 without phage addition. Results are presented as mean values ± SD from three independent experiments. Please note that the bars are smaller than the sizes of the symbols.

### 3.7. Resistance of vB-EcoS_57-3 virions to external conditions and urine environment

The sensitivity of vB-EcoS_57-3 to various external conditions, including various temperatures, pH conditions, solvents, and detergents was tested (**Table 3**). The virions appeared relatively resistant to freezing, pH values, or osmotic shock, although they could not survive under conditions of 65°C and a pH of 2. Resistance to disinfectants, organic solvents, and detergents differed depending on the nature of the tested compound and was the highest for ethanol and chloroform and the lowest for 0.1% cetyltrimethylammonium bromide (CTAB), in which phages lost their activity completely.

**Table 3.** Resistance of vB-EcoS_57-3 phage virion to physical and chemical agents. The values presented in the table represent the percentage of viable phage particles under the specified conditions.

| Phage name | Temperature |  |  | pH (1h, 37°C) |  |  | Osmotic shock | 0.1% CTAB (1 min) | 0.1% Sarkosyl (10 min) | 63% Ethanol (1h) | 50% DMSO (10 min) | Chloroform (1.5 h) |
| --- | --- | --- | --- | --- | --- | --- | --- | --- | --- | --- | --- | --- |
|  | -20°C (24h) | 40°C (40 min) | 62°C (40 min) | pH 2 (1h) | pH 4 (1h) | pH 10 (1h) |  |  |  |  |  |  |
| vB-EcoS_57-3 | 100 | 100 | 1.5 | 0 | 100 | 100 | 100 | 0 | 50 | 100 | 50 | 100 |

In artificial urine (**Figure 8A**), vB-EcoS_57-3 remained relatively stable across all tested pH values (4.5, 5.8, and 6.5), with only minor fluctuations in PFU counts over time. At pH 5.8 and 6.5, phage titers were consistently maintained at high levels throughout the incubation period, indicating favorable conditions for virion stability. In contrast, at pH 4.5, a slight reduction in PFU was observed at certain time points (4, 6, 8 h), suggesting that more acidic conditions may have a modest negative impact on phage stability. Nevertheless, even under acidic conditions, the decrease was not pronounced, demonstrating that vB-EcoS_57-3 retains antibacterial activity in artificial urine, significantly reducing bacterial counts up to 10% of the initial amount, after 24-hour of incubation (**Figure 8B**). In contrast, in the absence of phage particles, bacterial populations remained stable after 24 hours of incubation, with CFU values close to or only slightly reduced from the initial levels (**Figure 8C**). The survival of bacteria was correspondingly high, indicating that artificial urine at pH 4.5 alone does not significantly inhibit bacterial growth. This highlights that the observed reduction in bacterial counts in **Figure 8B** is specifically attributable to phage activity rather than environmental stress.

**Figure 8.**
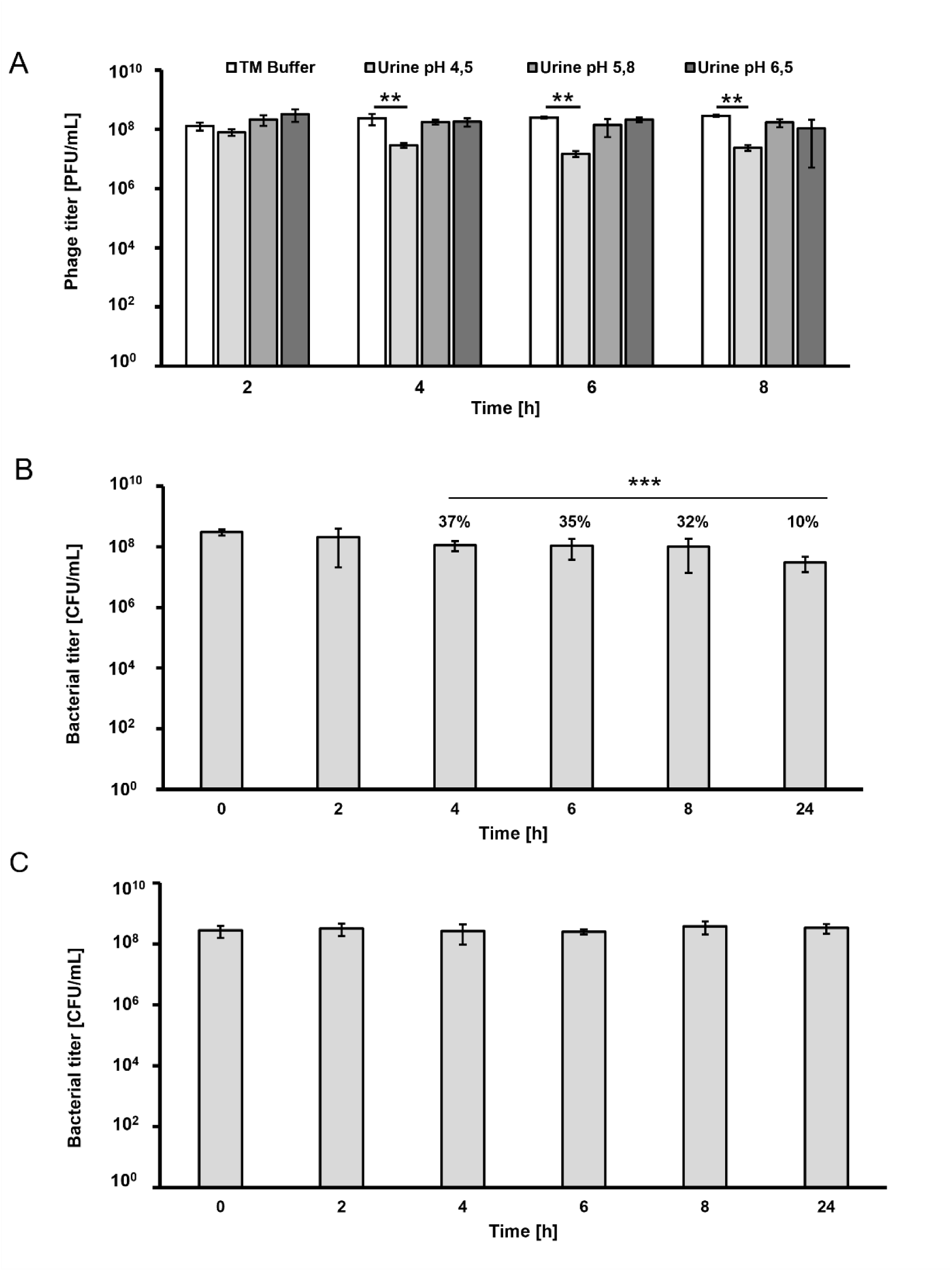
Stability and activity of phage vB-EcoS_57-3 in artificial urine. (**A**) Stability analysis of phage virions was performed at 2, 4, 6, and 8 hours of incubation in artificial urine with different pH (4.5; 5.8; 6.5). The number of phage particles was presented as the number of plaque-forming units (PFU) present in 1 mL of urine. (**B**, **C**) Activity of bacteriophage vB-EcoS_57-3 presented as the number of bacterial cells of the EC57 strain that survived 24-hour incubation in artificial urine at pH = 4.5, in the presence (**B**) and without the addition of phage particles (**C**). The number of bacterial cells was expressed as colony-forming units (CFU) per 1 mL of urine. The number of cells that survived the infection relative to the initial number of bacteria is expressed in percentages above the bars. The Student’s *t*-test was used to calculate statistical significance. Statistically significant results are marked with asterisk symbols: *P* < 0.01 (**) and *P* < 0.001 (***).

### 3.8. The safety of vB-EcoS_57-3 toward human cells

To test the safety of phage vB-EcoS_57-3 to mammalian cells, the viability of the human bladder cancer cell line, T24 (**Figure 9A**), and the human embryonic kidney line, HEK293 (**Figure 9B**), was assessed after treatment with the phage lysate at 3 concentrations (10^10^, 10^9^, and 10^8^ PFU/mL). The endotoxin level in the applied phage lysate was estimated at 0.57 ± 0.03 EU/mL, which is considered to be safe for mammalian cells. Besides, the stability of phage particles was tested after 24-hour incubation with mammalian cells T24 (**Figure 9C**) and HEK293 (**Figure 9D**). The addition of phages to T24 or HEK293 cultures resulted in no significant changes detected in the number of viable cells relative to the negative control, which included non-treated cultures with viability estimated at the level of 100% (**Figure 9A, B**). The lack of a drop in cell survival below 70% in all tested cases (except the positive control) suggested no deleterious effect of the tested phage on the viability of mammalian cells.

**Figure 9.**
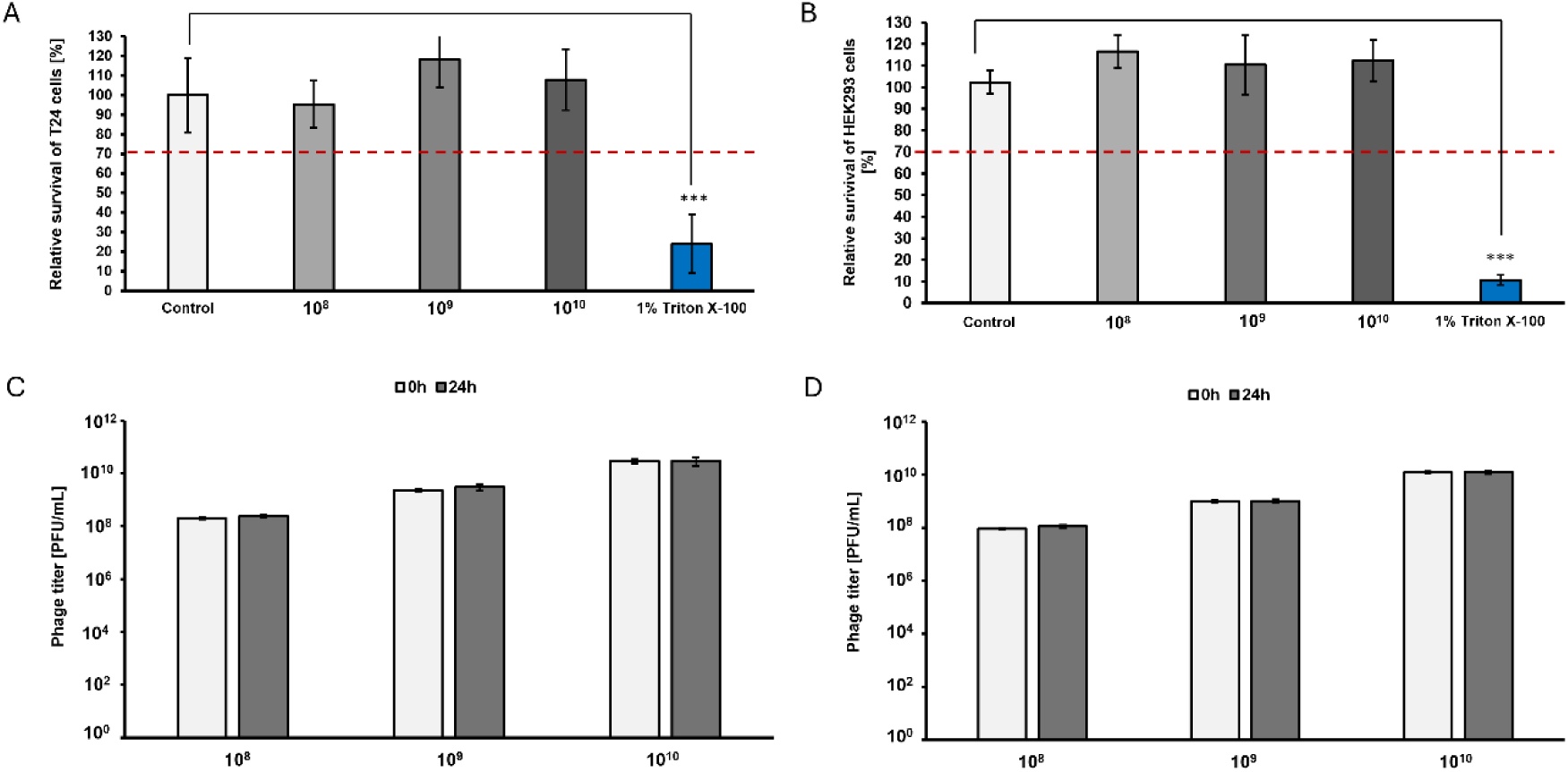
Assessment of cytotoxicity of vB-EcoS_57-3 phage particles to mammalian cells T24 (**A**) and HEK293 (**B**) after 24 h incubation. The phage lysate tested at 3 concentrations (10^10^, 10^9^, and 10^8^ PFU/mL). The letter T means a drop in cell survival below the red dotted threshold line at 70%, which qualifies the agent as toxic. The stability of phage vB-EcoS_57-3 during incubation with mammalian cells T24 (**C**) and HEK293 (**D**) shown as the number of phage particles in 1 mL (PFU/mL) of lysate measured at time 0 and after 24 hours of incubation. The Student’s *t*-test was used to calculate statistical significance, which are marked with asterisk symbols: *P* < 0.001 (***).

### 3.9. In silico analysis of the 57_3Lys endolysin properties

The vB-EcoS_57-3 genome analysis allowed for the identification of the endolysin (GenBank accession number YEN46648.1) in the predicted lysis cassette. *In silico* analysis (**Table 4**) revealed that the analyzed endolysin gene is located between nucleotides 40,824 and 41,309 in the phage genome and has a length of 486 bp, encoding a relatively small protein of 161 amino acids. The compact size of both the gene and the encoded protein is characteristic of phage lytic enzymes, particularly endolysins. The predicted molecular weight of 17.55 kDa further supported the classification of this protein as a small enzymatic protein, consistent with known phage lysins. The isoelectric point (pI) of 9.7 indicates that the protein is basic in nature, suggesting a net positive charge at physiological pH. This property may facilitate interactions with negatively charged bacterial cell wall components, such as peptidoglycan.

**Table 4.**
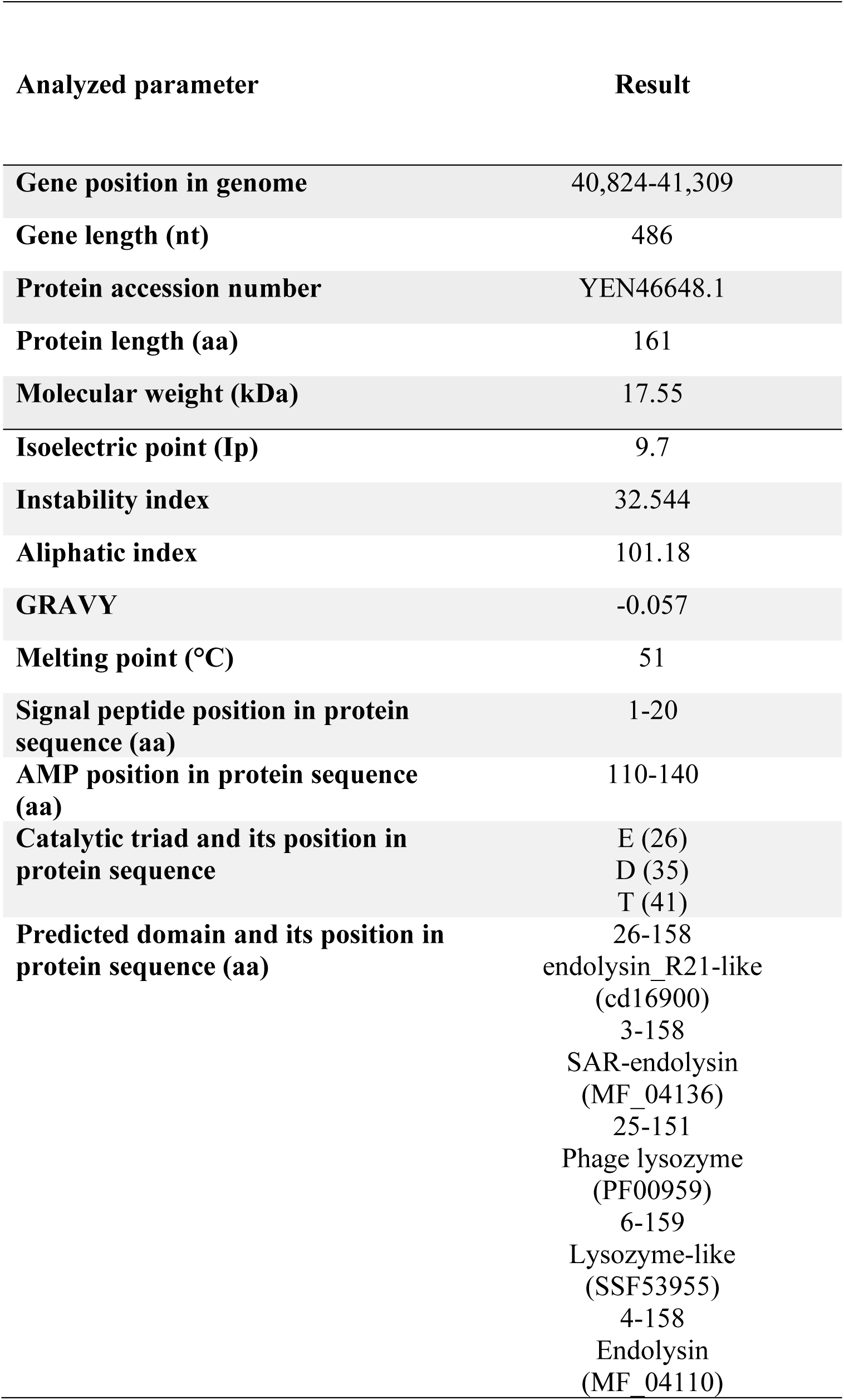
Characteristics of the 57_3Lys endolysin.

The instability index of 32.544 classifies the protein as stable (values below 40 indicate stability), suggesting that it is likely to maintain its structural integrity under standard conditions. The aliphatic index of 101.18 is relatively high, indicating a substantial proportion of aliphatic amino acids, which is typically associated with enhanced thermal stability. The GRAVY score of -0.057 indicates that the protein is overall slightly hydrophilic. This suggests good solubility in aqueous environments, which is advantageous for enzymes that act on cell wall substrates in extracellular or periplasmic spaces. Domain prediction analysis identified multiple overlapping domains associated with lysozyme and endolysin activity, including SAR endolysin and lysozyme domains at positions 3-158 and 25-151, respectively (**Table 4**), thus confirming that the protein belongs to the phage endolysin/lysozyme family, responsible for degrading bacterial peptidoglycan during host cell lysis.

### 3.10. Identification of motifs characteristic for class II SAR endolysins within 57_3Lys

Searching for motifs characteristic for phage endolysins revealed three important domains within the 57_3Lys organization structure, typical for class II SAR endolysins (**Figure 10**), represented by R_21_, the endolysin of the lambdoid phage 21 (Xu et al., 2004; Kuty et al., 2010). These enzymes have the canonical E-D-T catalytic triad and no C residue in the SAR domain.

**Figure 10.**
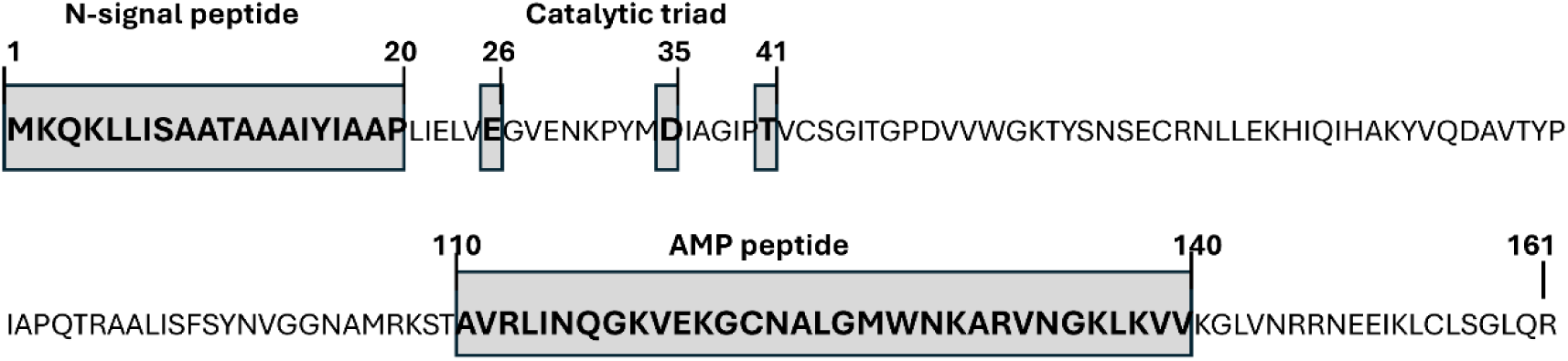
Sequence of 57_3Lys endolysin with predicted motifs characteristic for class II SAR endolysins: a typical signal peptide with features of a SAR domain, including a moderately hydrophobic segment encompassing the first 20 amino acids; putative catalytic residues (E26, D35, and T41); and an AMP peptide at the C-terminus, between 110 and 140 amino acids.

The N-terminal region (residues 1-20) of 57_3Lys contains a clearly identifiable signal peptide with SAR-domain properties, as indicated by the sequence MKQKLLISAATAAAIYIAAP (**Figure 10** and **Supplementary Figure S2**). This segment is moderately hydrophobic, has no cysteine residue, and is predicted to function as a transient membrane anchor. Its identification by SignalP 6.0 suggests that the 57_3Lys protein, as a typical class II SAR endolysin, is targeted to the Sec-dependent secretory pathway, where it is translocated across the cytoplasmic membrane and remains anchored within the membrane by the N-terminal signal peptide until membrane depolymerization occurs. Unlike classical signal peptides, the SAR domain of class II endolysins is not immediately released but instead retains the enzyme in an inactive, membrane-tethered state.

Downstream of the signal peptide, the 57_3Lys protein sequence forms the catalytic core of the endolysin as revealed by NCBI Conserved Domain software (**Figure 10** and **Supplementary Figure S2**). Within this region, three key residues, E26, D35, and T41, are highlighted as the putative catalytic motif. This arrangement corresponds to the conserved E-(X)₈-D-(X)₅-T motif, which is characteristic for class II SAR endolysins rather than the canonical cysteine-dependent catalytic triads of class I SAR endolysins. The positioning of these residues suggests a hydrolytic mechanism targeting peptidoglycan, although enzymatic activity is dependent on conformational rearrangements that reposition the catalytic glutamate upon release from the membrane, rather than solely on residue composition.

In addition, the CAMP analysis predicted an antimicrobial peptide (AMP) region between residues 110-140, located within the C-terminal part of the 57_3Lys protein (**Figure 10** and **Supplementary Figure S2**). This region shows strong similarity to cecropin-like α-helical antimicrobial peptides, particularly in its enrichment in positively charged and hydrophobic residues. Despite relatively low primary sequence identity (∼20-25%) to Cecropin A from *Hyalophora cecropia*, the physicochemical properties, such as cationic charge distribution and amphipathic character, are conserved, suggesting a similar membrane-disruptive mode of action. This implies that the 57_3Lys protein may possess a dual function: enzymatic degradation of the peptidoglycan and direct disruption of bacterial membranes.

Phylogenetic analysis of 20 orthologous endolysins revealed that the endolysin of vB-EcoS_57-3 (YEN46648.1) clustered with endolysins from related phages (**Figure 11**). The vB-EcoS_57-3 endolysin of vB-EcoS_57-3 formed a well-supported sister group with the endolysin of vB_EcoS_ACG-M12 (SH-aLRT/UFBoot: 85.6/92), while the endolysin of vB_EcoS-12210I was positioned as the next related lineage. The analyzed endolysins generally showed similar physicochemical properties and domain composition (**Supplementary Table S3**). Most were 160-161 amino acids long, with predicted molecular weights of approximately 17.3-17.6 kDa and basic isoelectric points.

**Figure 11.**
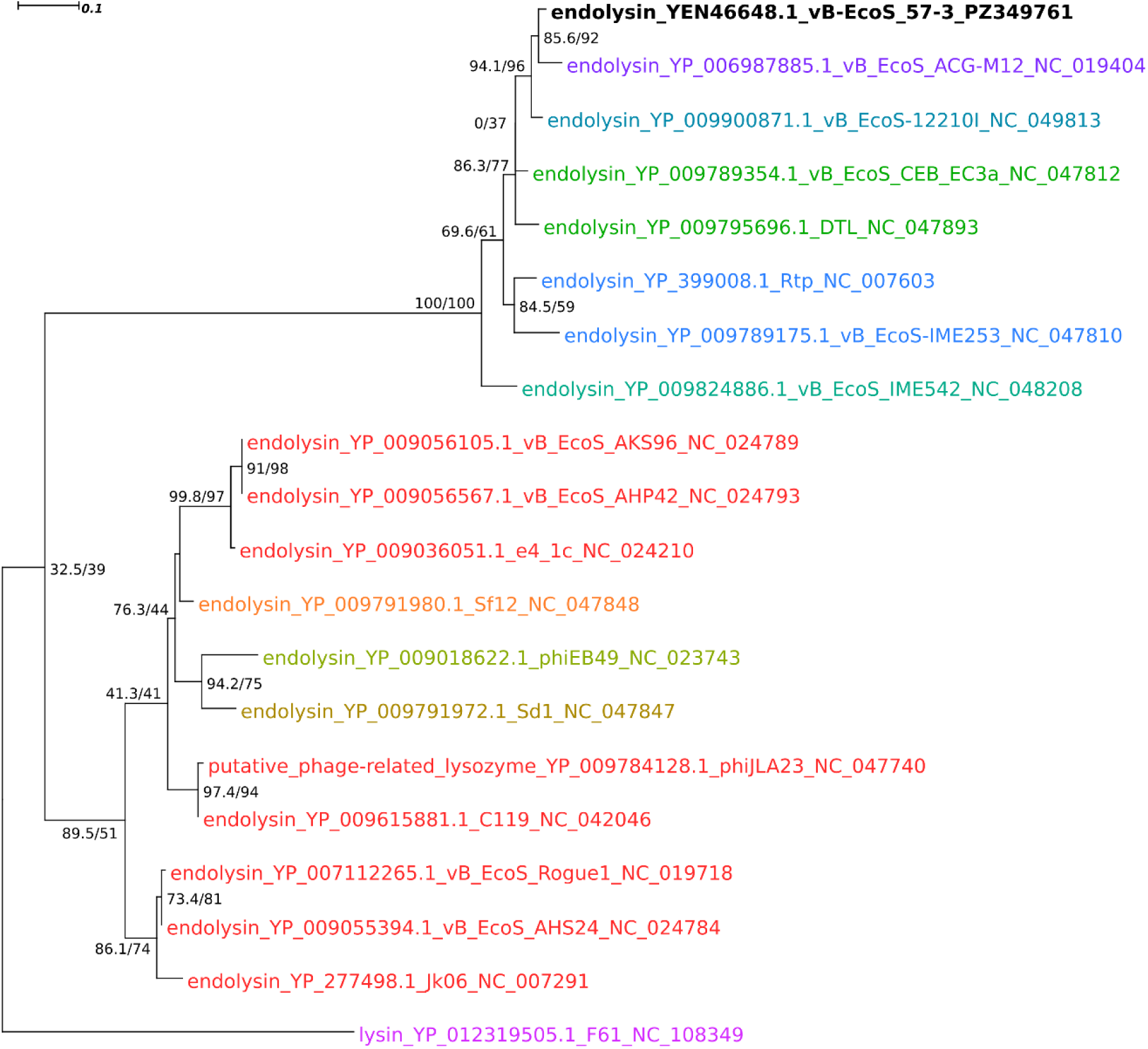
Phylogenetic relationships of the endolysin from bacteriophage vB-EcoS_57-3 and orthologous endolysins from related phages. The maximum-likelihood phylogenetic tree was constructed based on amino acid sequences of 20 orthologous endolysins. Branch support values at the nodes are presented as SH-aLRT support (%) / ultrafast bootstrap support (%), based on 1,000 replicates each. The endolysin of vB-EcoS_57-3 (YEN46648.1) is highlighted in bold. Different protein name colors indicate the genus-level taxonomic assignment of the analyzed phages as follows: red *Rogunavirus*, orange *Eastlansingvirus*, dark yellow *Wilsonroadvirus*, bottle green *Lindendrivevirus*, green *Loudonvirus*, teal *Christensenvirus*, dark cyan *Veterinaerplatzvirus*, blue *Rtpvirus*, violet *Guelphvirus*, pink *Tlsvirus*. The scale bar represents the number of amino acid substitutions per site.

Domain analysis further demonstrated a highly conserved architecture among the analyzed endolysins. The vB-EcoS_57-3 endolysin contained signatures corresponding to phage lysozyme (PF00959), endolysin_R21-like (cd16900), lysozyme-like (SSF53955), endolysin (MF_04110), and SAR-endolysin (MF_04136), a domain profile shared by most of the analyzed proteins (**Table 4**, **Supplementary Table S3**). Notable exceptions included the shorter putative phage-related lysozyme of phiJLA23 (131 aa), for which the SAR-endolysin signature was not detected. Overall, the phylogenetic and domain analyses indicate that the vB-EcoS_57-3 endolysin belongs to a group of closely related, structurally conserved phage lysozymes.

### 3.11. Involvement of the Sec machinery in 57_3Lys-mediated lysis of E. coli bacteria

To examine whether the identified SAR motif functions as a secretion sequence for Sec translocase and whether Sec machinery is responsible for 57_3Lys secretion, the endolysin was produced from the plasmid in the presence of NaN_3_ (**Figure 12**), a well-known inhibitor of *E. coli* Sec translocase (Bai et al., 2020). In this experiment, the host cell lysis kinetics of 57_3Lys were determined by adding various concentrations of NaN_3_. Increasing concentrations of NaN_3_ inhibited the cell lysis process induced by the 57_3Lys produced from the plasmid after the addition of IPTG and arabinose (**Figure 12**). The addition of 5 mM NaN_3_ reduced the lytic effect of the 57_3Lys endolysins by about half an OD_600_ unit, whereas the addition of 10 mM NaN_3_ almost completely inhibited it and returned the culture turbidity to the level of the variant with non-produced 57_3Lys. This line of evidence implies that the Sec system might be involved in 57_3Lys-mediated cell lysis *in vitro*.

**Figure 12.**
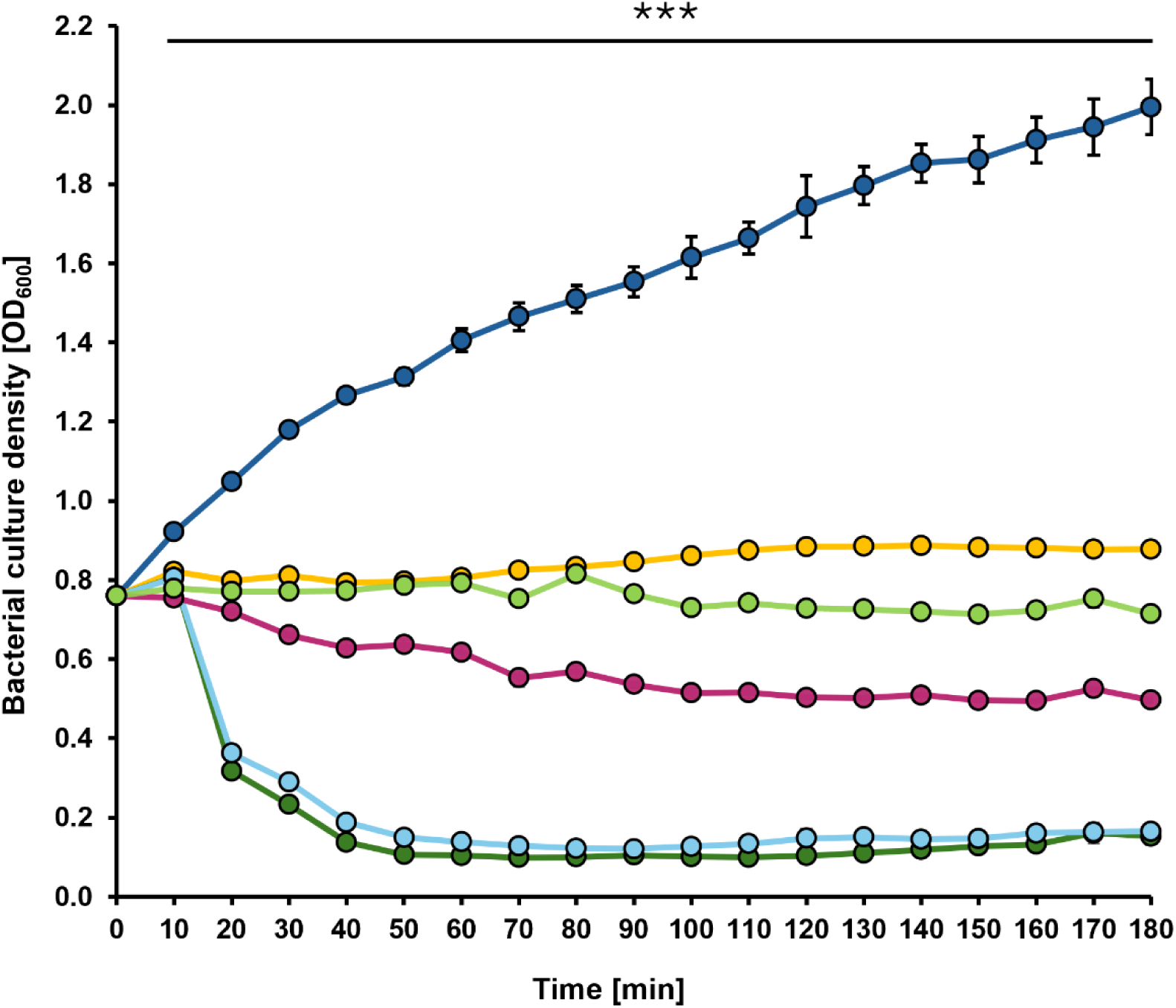
Kinetics of *E. coli* cell lysis by the 57_3Lys endolysin in the presence of the Sec translocase inhibitor, NaN_3_. In all experiments, expression of 57_3Lys from plasmid in *E. coli* cells was induced by adding 0.5 mM of IPTG and 0.2% arabinose, except for the controls. Dark blue circles indicate samples without NaN₃, IPTG and arabinose (control 1), orange circles indicate samples treated with 10 mM NaN₃ in the absence of IPTG and arabinose (control 2), while dark green, light blue, dark pink, and light green circles represent samples treated with 0, 1, 5, and 10 mM NaN₃, respectively. All experiments for the cell lysis kinetics were performed in triplicate and the results of each treatment are represented by the mean ± SD. Statistical analyses were performed by the Student’s *t*-test. The significance of differences between control 1 and particular variants are observed from the 10^th^ minute and marked by asterisks: *P* < 0.001 (***). Please note that the bars are smaller than the sizes of the symbols.

### 3.12. Functional analysis of the purified 57_3Lys lytic activity

Functional assays showed that overexpression of 57_3Lys significantly reduced bacterial turbidity, indicating effective lysis (**Figure 12**), so we decided to purify this protein and analyze its lytic action against the EC57 clinical UPEC isolate. The Ni^2+^-NTA affinity chromatography allowed us to obtain a fraction of the purified 57_3Lys endolysin corresponding to the predicted molecular weight of approximately 17.55 kDa, further verified by mass spectrometry (**Supplementary Figure S3**).

Purified 57_3Lys was used in turbidity and cell number reduction assays that demonstrated its antibacterial activity, significantly reducing the OD_600_ value of the EC57 bacterial culture and the amount of viable EC57 cells expressed as CFU/mL (**Figure 13 A**, **B**). Importantly, the antibacterial effect was observed after 24 h incubation of the bacterial cells with the 57_3Lys endolysin and was not detected during the first 60 minutes of the treatment (**Supplementary Figure S4**). Curiously, incubation with the commercially purchased lysozyme has no significant antibacterial effect, probably due to a lack of access of the lysozyme to the peptidoglycan layer. The observed antibacterial effect of the 57_3Lys in this case may be the result of the outer membrane destabilization by the internal AMP peptide action.

**Figure 13.**
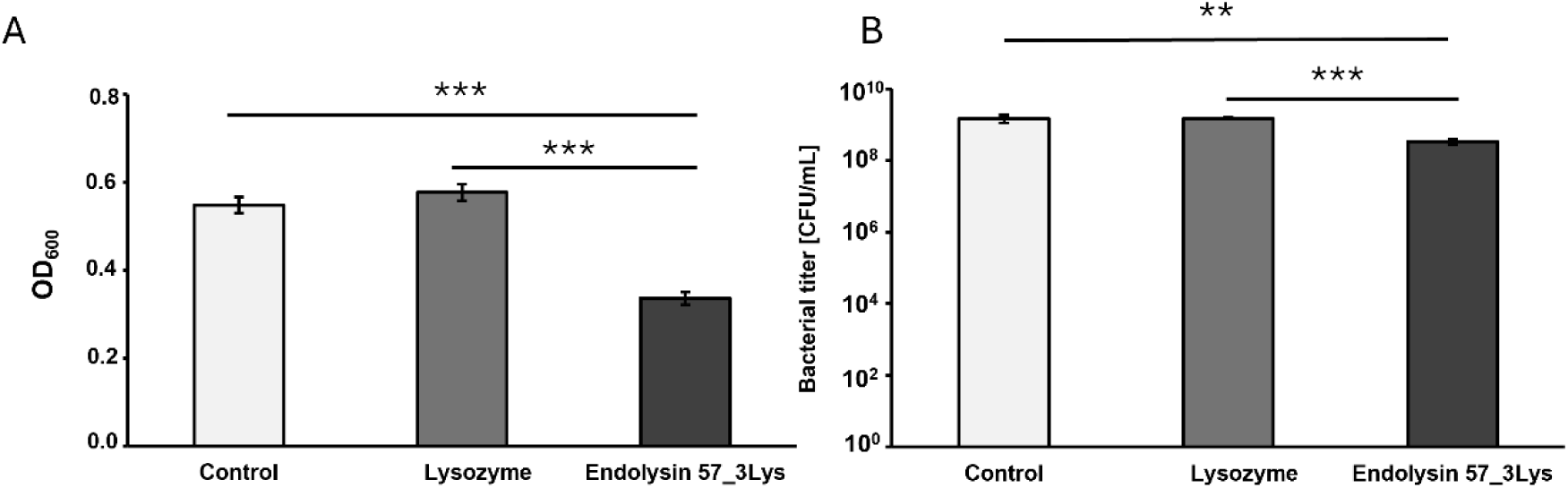
Turbidity (**A**) and cell number (**B**) reduction assays of EC57 culture by purified 57_3Lys endolysin after 24 hours of incubation. Each experiment was performed in triplicate, and the results of each treatment are represented by the mean ± SD. Statistical analyses were performed by the Student’s *t*-test. The significance of differences between endolysin variant and particular control variants are observed and marked by asterisks: *P* < 0.01 (**) and *P* < 0.001 (***).

## 4. Discussion

In this study, we characterized the newly isolated UPEC-infecting bacteriophage vB-EcoS_57-3 and its endolysin 57_3Lys. The phage efficiently lysed EC57, showed activity against selected clinical UPEC isolates, including antibiotic-resistant strains, remained active under urinary conditions, and did not measurably affect the viability of tested human bladder and kidney cell lines. 57_3Lys was identified as a class II SAR endolysin, and functional data indicated that its lytic activity depends strongly on the host Sec pathway. Importantly, purified 57_3Lys also reduced the turbidity and the number of viable EC57 cells without an externally added outer-membrane permeabilizer, suggesting intrinsic activity against Gram-negative bacteria.

Sequence analysis identified an N-terminal, moderately hydrophobic segment of approximately 20 amino acids with properties expected for a SAR domain, together with the E(X)8-D(X)5-T catalytic motif characteristic of class II SAR endolysins. This architecture differs from canonical cytoplasmic phage endolysins, whose access to the periplasm generally depends on holin-generated membrane lesions. In SAR systems, the N-terminal signal-anchor engages the host Sec machinery, allowing translocation of most of the endolysin through the cytoplasmic membrane while the SAR domain remains transiently tethered to it. Membrane depolarization subsequently releases the N-terminal signal and permits formation of the active periplasmic enzyme. The paradigm was established for the P1 endolysin Lyz, for which Sec-dependent export and non-proteolytic release of the N-terminal SAR sequence were demonstrated experimentally (Xu et al., 2004). Related pinholin-SAR systems have been described for phage 21 and phiKMV, illustrating that membrane depolarization rather than formation of large protein-conducting holes can control the timing of lysis (Park et al., 2007; Briers et al., 2011).

Our functional data obtained for 57_3Lys support this pathway. Increasing concentrations of sodium azide, an inhibitor of the *E. coli* SecA-dependent translocation machinery (Bai et al., 2020), progressively suppressed lysis following induction of 57_3Lys expression. At 5 mM NaN_3_ the lytic effect was markedly reduced, whereas 10 mM NaN_3_ almost completely abolished the decrease in culture turbidity, bringing the phenotype close to that of the variant without induced 57-3Lys production. Because sodium azide can exert cellular effects beyond a single protein-transport event, these experiments should not be interpreted as direct visualization of 57_3Lys translocation. Nevertheless, the dose-dependent inhibition, considered together with the predicted N-terminal SAR signal and phylogenetic placement of 57_3Lys among class II SAR endolysins, provides convergent functional evidence that Sec-mediated export is required for efficient 57_3Lys-mediated lysis.

The absence of cysteine in the SAR region of 57_3Lys further supports a class II SAR endolysin-type activation model. In contrast to class I SAR endolysin P1 Lyz, controlled by topological, conformational and disulfide-dependent changes after membrane release (Xu et al., 2005), class II-like enzymes lacking the cysteine residues within the SAR domain are expected to rely primarily on conformational repositioning of catalytic residues E, D, T after release from the membrane (Woźnica et al., 2015). Importantly, the described 57_3Lys features also argue against describing its N-terminal region as a conventional cleavable signal peptide. In the established class II SAR paradigm, the signal-anchor is retained rather than removed by signal peptidase and is released from the membrane upon depolarization (Xu et al., 2004; Park et al., 2007).

A next particularly notable finding was the antibacterial activity of externally applied 57_3Lys against intact EC57 cells. Because the Gram-negative outer membrane normally restricts access of endolysins to peptidoglycan, this activity suggests that 57_3Lys possesses features facilitating interaction with or passage across the outer membrane. The effect developed slowly, becoming significant after 24 h rather than within the first 60 min, indicating that outer-membrane access may represent the rate-limiting step. Similar intrinsic activity has been reported for ABgp46, LysMK34, LysAm24, LysECD7 and LysSi3 (Oliveira et al., 2016; Antonova et al., 2019; Abdelkader et al., 2022). More recently, PA16cLys demonstrated activity against mature biofilms of uropathogenic *P. aeruginosa* without an additional permeabilizer, further supporting the potential of intrinsically active lysins in urinary infections (Zhang et al., 2026).

The predicted C-terminal AMP-like region of 57_3Lys, enriched in positively charged and hydrophobic residues, together with the high predicted pI (∼9.7), provides a plausible explanation for this activity. Such cationic and amphipathic regions may interact electrostatically with lipopolysaccharide and destabilize the outer membrane. Similar principles underlie cecropin A-fused lysins and artilysins (Briers et al., 2014; Abdelkader et al., 2022). However, the contribution of this region to 57_3Lys activity remains to be experimentally demonstrated. The relatively slow killing compared with ABgp46 and engineered membrane-active lysins such as Art-175 (Briers et al., 2014; Oliveira et al., 2016) suggests that improving outer-membrane penetration could substantially enhance 57_3Lys antibacterial efficacy.

Finally, the present study demonstrates that both the parental phage vB-EcoS_57-3 and its endolysin 57_3Lys possess features relevant to this approach. The phage provides self-amplifying, receptor-dependent killing and showed stability and antibacterial activity in artificial urine, whereas the purified endolysin bypasses the need for phage adsorption and replication. This distinction may be particularly relevant when phage resistance emerges. A recent report on PA16cLys is illustrative, as the endolysin retained activity against phage-resistant *P. aeruginosa* mutants (Zhang et al., 2026). Overall, the findings support further investigation of both vB-EcoS_57-3 and 57_3Lys as potential alternative or adjunctive strategies against antibiotic-resistant UPEC, while highlighting the need to improve and mechanistically define the extracellular activity of 57_3Lys. Besides, combined or sequential phage–endolysin treatment could be investigated as a potential strategy to improve antibacterial efficacy and address phage resistance.

## CRediT authorship contribution statement

**Wojciech Wesołowski:** Writing – original draft, Visualization, Investigation, Methodology, Data curation

**Sylwia Bloch:** Methodology, Investigation, Writing – review & editing

**Grzegorz Czerwonka:** Methodology, Investigation, Writing – original draft

**Ernest Jagieła:** Investigation, Visualization

**Łukasz Grabowski:** Investigation, Visualization

**Marta Jeschke:** Investigation, Visualization

**Jakub Jacewicz:** Investigation, Visualization

**Emilia Piwnicka:** Writing – original draft, Data curation

**Joanna Morcinek-Orłowska:** Methodology, Data curation

**Hanna Loika:** Investigation, Visualization

**Paulina Czaplewska:** Investigation, Methodology

**Grzegorz Węgrzyn:** Writing – review & editing, Formal analysis.

**Wioletta Adamus-Białek:** Writing – review & editing, Data curation, Supervision

**Aleksandra Łukasiak**: Investigation, Funding acquisition, Data curation.

**Bożena Nejman-Faleńczyk:** Writing – original draft, Writing – review & editing, Supervision, Formal analysis, Conceptualization.

## Supplementary materials

**Supplementary Figure S1.**
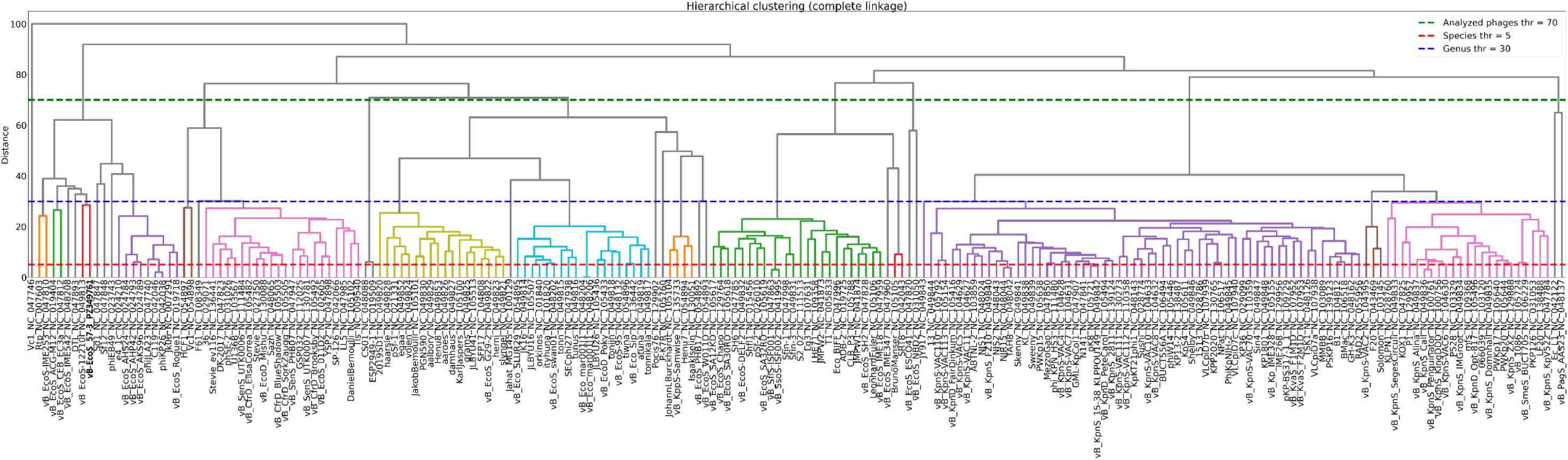
VIRIDIC-based clustering of phage vB-EcoS_57-3 and reference phages from the family *Drexlerviridae*. Hierarchical clustering based on pairwise intergenomic distances calculated using VIRIDIC for phage vB-EcoS_57-3 and reference phages from the family *Drexlerviridae*. Clustering was performed using complete linkage. Dashed lines indicate the distance thresholds corresponding to the species level (5%; red), genus level (30%; blue), and for limiting the number of the analyzed phages up to 20 cases (70%; green).

**Supplementary Figure S2.**
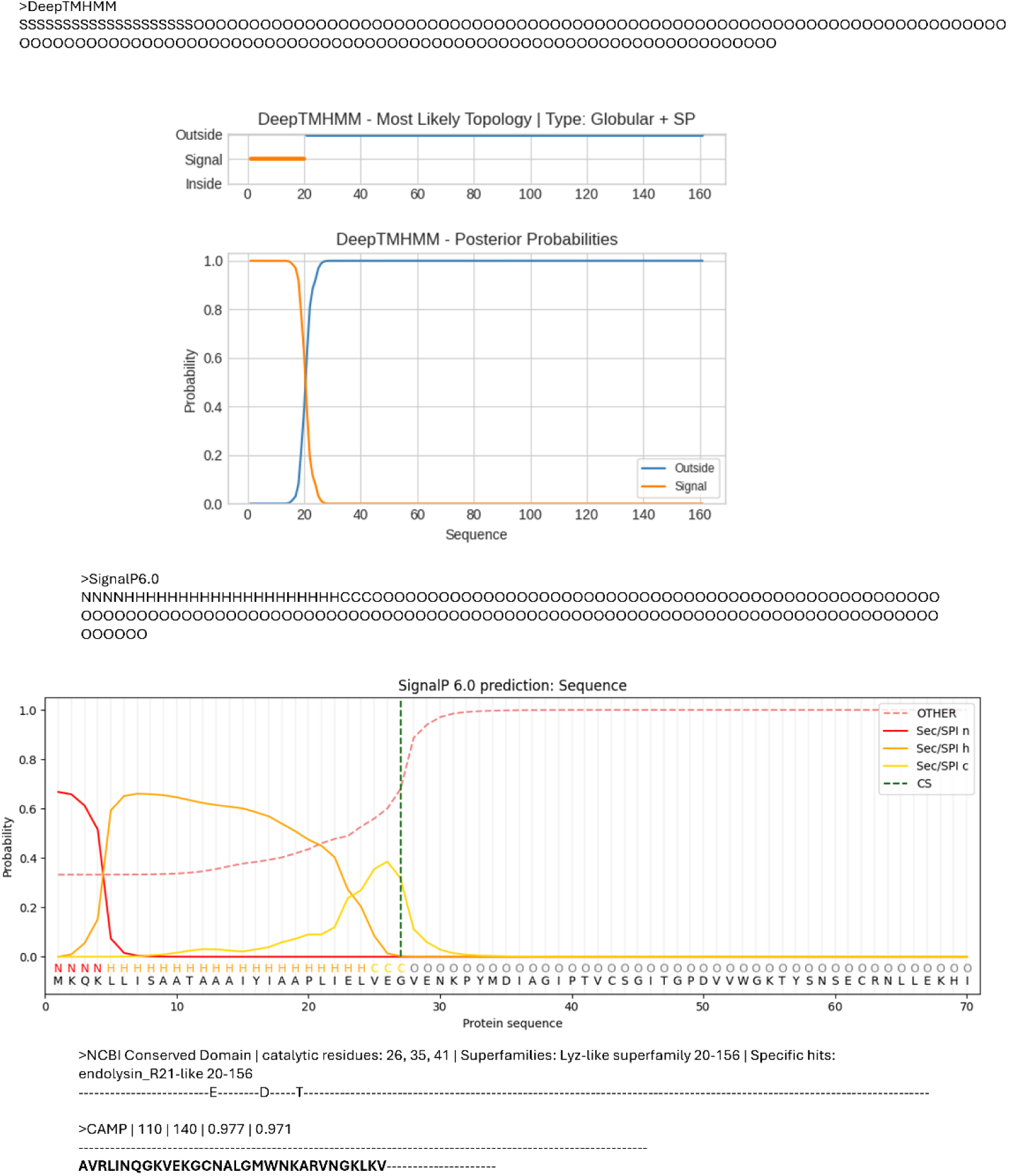
Results of bioinformatic analysis of the 57_3Lys endolysin sequence confirming the presence of a 20 aa-long signal molecule at the N end (>SignalP 6.0 and >Deep TMHMM); a catalytic triad E-D-T encompassing 26^th^, 35^th^, and 41^st^ residues, respectively (>NCBI Conserved Domain); and an AMP peptide occurring at a position between 110 and 140 amino acids (>CAMP).

**Supplementary Figure S3.**
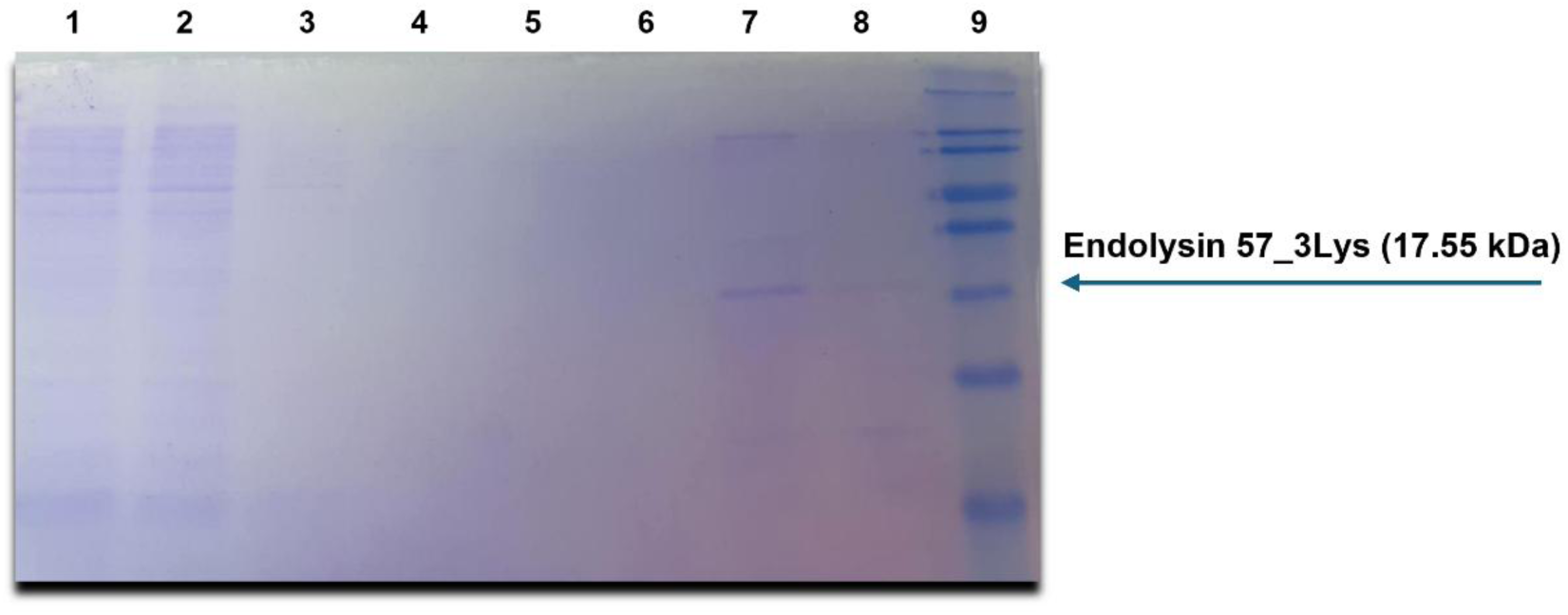
SDS-PAGE analysis of protein purification fractions obtained after Ni2+-NTA affinity chromatography of 57_3Lys endolysin. The 7^th^ lane, corresponding to the 2^nd^ elution fraction contains a prominent band at approximately 17.55 kDa, corresponding to the predicted molecular weight of 57_3Lys, indicating successful expression and purification. The remaining gel lanes contain: 1 - Lysate of *E. coli* BL21-AI [pET26b_57_3Lys]; 2 - Flow through; 3 - Wash 1; 4 - Wash 2; 5 - Wash 3; 6 - 1^st^ elution fraction; 8 - 3^rd^ elution fraction. The last, 9^th^ lane contains SeeBlue® Pre-Stained Protein Standard marker (kDa). The gel was stained with BlueStain Sensitive Plus.

**Supplementary Figure S4.**
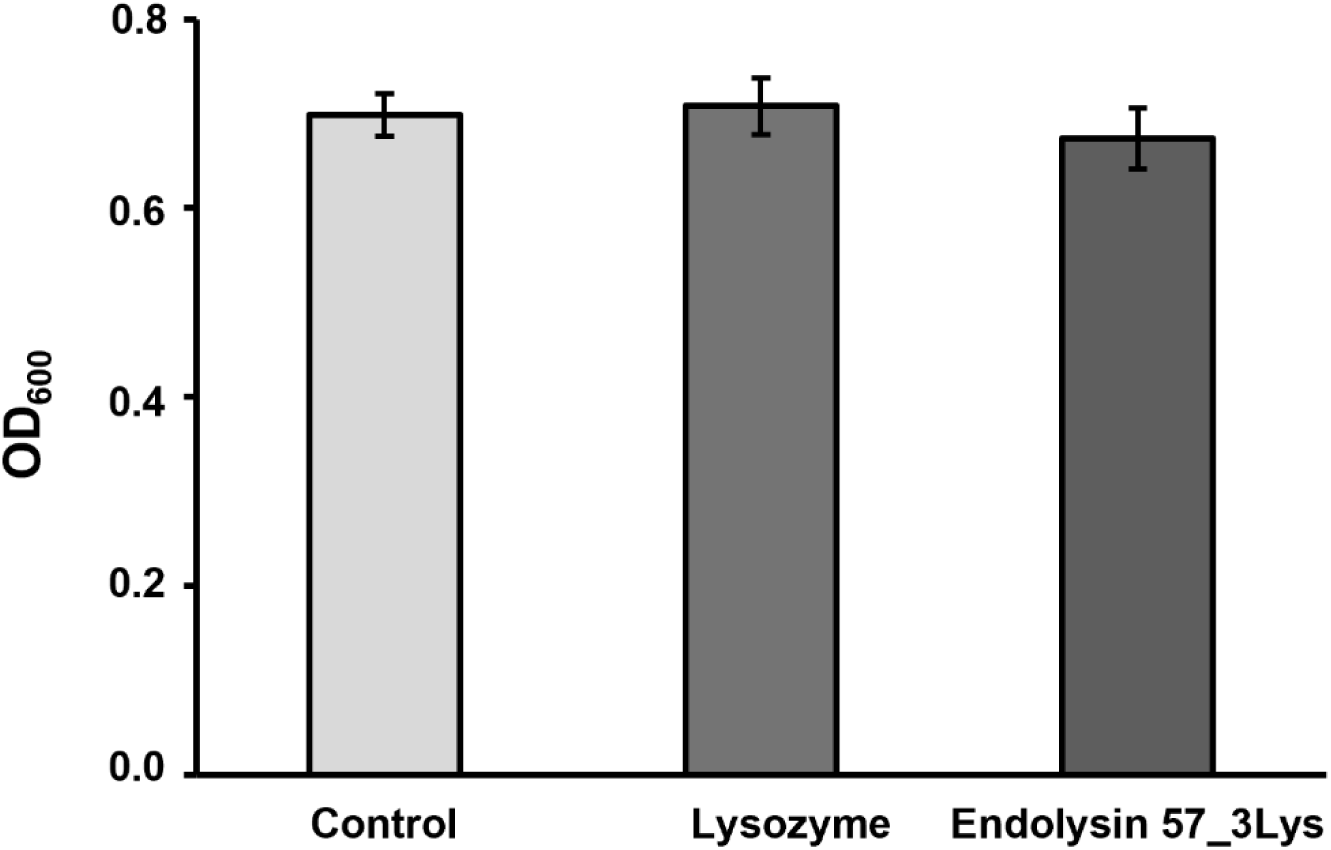
Turbidity reduction assays of EC57 culture by purified 57_3Lys endolysin after 60 minutes of incubation. Turbidity reduction assays were performed in triplicate and the results of each treatment are represented by the mean ± SD.

**Supplementary Table S1. Intergenomic distances among phage vB-EcoS_57-3 and reference phages calculated using VIRIDIC.** Pairwise intergenomic distances between phage vB-EcoS_57-3 and reference phages from the family *Drexlerviridae*, calculated using VIRIDIC. Values are expressed as genomic distances (%), where lower values indicate higher intergenomic similarity.

**Supplementary Table S2.** Orthologous protein groups used for phylogenetic analysis of vB-EcoS_57-3. The first six columns provide information on the alignments and their use in the phylogenetic analysis. Alignment length - length of the protein sequence alignment (number of amino acid positions); Parsimony - number of parsimony-informative sites in the alignment; Singleton - number of singleton sites in the alignment; Constant – number of constant sites in the alignment; Included in phylogeny - indicates whether the corresponding orthologous protein group was included in the phylogenetic analysis, together with the justification for its inclusion (+) or exclusion (-), where applicable; Evolution model - best-fit amino acid substitution model selected for the corresponding alignment partition by ModelFinder Plus implemented in IQ-TREE. The remaining columns correspond to individual phages, with each column header providing the phage name and its GenBank accession number. Cells in these columns contain the annotated protein assigned to the corresponding orthologous group and its protein accession number.

**Supplementary Table S3.** Physicochemical properties and functional annotation of the endolysin from bacteriophage vB-EcoS_57-3 and orthologous endolysins included in the phylogenetic analysis. The table summarizes the predicted physicochemical properties and functional annotations of the 20 orthologous endolysins included in the phylogenetic analysis (Figure 11). For each protein, the phage name, protein accession number, protein length, predicted molecular weight (MW), theoretical isoelectric point (pI), instability index, aliphatic index, and GRAVY value are provided. Functional annotations and sequence signatures identified using InterPro are also presented, including phage lysozyme, endolysin_R21-like, lysozyme-like, endolysin, and SAR-endolysin signatures, if detected.

